# Tasselyzer, a machine learning method to quantify maize anther exertion, based on PlantCV

**DOI:** 10.1101/2021.09.27.461799

**Authors:** Chong Teng, Noah Fahlgren, Blake C. Meyers

## Abstract

Male fertility in maize involves complex genetic programming affected by environmental factors. Evaluating the presence and proportion of fertile anthers is crucial for agronomic purposes. Anthers in maize emerge from male-only florets, and quantifying anther exertion is a key indicator of male fertility; however, traditional manual scoring methods are subjective. To address this limitation, we developed an automated method, *Tasselyzer*, for large-scale analysis. This image-based program uses the PlantCV platform to provide a quantitative assessment of anther exertion, capturing regional differences within the tassel based on the distinct color of anthers. We successfully applied this method to diverse maize lines to demonstrate its utility for research and breeding programs.

**Significance Statement:** Tasselyzer is a novel image-based segmentation tool for automated, large-scale measurement of anther exertion and the impact of genetic and environmental variation on male fertility in maize.

## Introduction

Male fertility in maize is controlled by complex genetic programming involving many developmental steps. Normal development can be negatively impacted by environmental factors such as light, temperature, water, nutrient availability, and mechanical stress such as wind or storm. The ability to regulate male fertility has substantial agronomic utility; consequently, the ability to evaluate the presence and proportion of fertile anthers is a beneficial technique. Maize anthers emerge from male-only florets, with two florets – an upper and a lower floret in each spikelet – and hundreds of spikelets arrayed on a terminal tassel, well-separated from the female ear. The stamen is a compound organ with a short filament subtending each developing anther. On the day of anthesis, the filament elongates substantially to push the anther through the enclosing spikelet glumes into the air, which initiates pollen shed. The filaments accompanying sterile anthers rarely elongate sufficiently to move the anther out of the spikelet (Egger and Walbot, 2015). Therefore, quantification of anther exertion is a useful method for scoring the extent of male fertility.

Traditionally, the scoring of anther presence is done manually using a simple scale from 0 = no anther exertion to 5 = full anther exertion (Yadava *et al*., 2021), however, manual scoring is subjective and varies among personnel evaluating the same tassels. Additionally, if there are regions within the tassel with a distinctive or inconsistent phenotype, this information is not captured; such regions could reflect suppression of anther development for one or a few days. To address these limitations, we developed ‘Tasselyzer’, an automated, image-based color trait analysis tool for maize tassel image segmentation built on the PlantCV naïve Bayes classifier (Fahlgren *et al*., 2015; Abbasi and Fahlgren, 2016; Gehan *et al*., 2017; Schuhl *et al*., 2022). Tasselyzer identifies anthers in two-dimensional tassel images based on distinct anther colors, separates them from other tassel parts, and quantifies the number of pixels of anthers separately from the other tassel parts, to generate an anther ratio. Then, Tasselyzer mimics human eye perception and calculates anther pixel ratios to evaluate the degree of exertion, thus quantifying the degree of male fertility in a maize plant. In a previous study, Tasselyzer enabled estimates of male fertility of individual tassels by measuring the ratios of anther pixels to those of the whole tassel, then pinpointed pheno-critical days of temperature treatments in the *dcl5* mutant (Teng *et al*., 2020).

In this study, we describe Tasselyzer and we examined how well Tasselyzer measures anther ratios compared to manually measuring the width of the main spike, which is a traditional method for tracking the anthesis process. Our goal was to determine if results from Tasselyzer are comparable to those obtained through traditional measurements. Additionally, we tested Tasselyzer’s effectiveness across a diverse set of maize lines, including 20 Nested Association Mapping (NAM) founder lines, to assess its applicability beyond specific genetic backgrounds. Finally, we discuss the reasons for using Tasselyzer and its potential for wider applications in research.

## Results

### Tasselyzer is a color-based tool for maize tassel image analysis

We captured 1,438 images of freshly detached tassels from 20 field-grown NAM founder lines and 2 inbred lines grown in greenhouses. These images were taken in an enclosed LED light box, ensuring consistent photo parameters (details provided in Materials and Methods and Supporting Table 1).

Tasselyzer segments images into multiple classes based on colors including: 1) dark non-plant background; 2) anthers in yellow or pink; 3) green parts of tassel, such as glumes, stems of the tassel, and sometimes palea. We selected 5 tassel images to create parameters, called the probability distribution function (PDF_1), used by Tasselyzer to analyze the remaining images, performing tasks including segmentation, generating masks for anthers and other tassel parts, and calculating anther ratios (Figure 1a-b). It is worth noting that an anther ratio is a measurement of a two-dimensional full or partial tassel image, but not an actual count of the anther numbers (details in Discussion).

**Figure 1.**
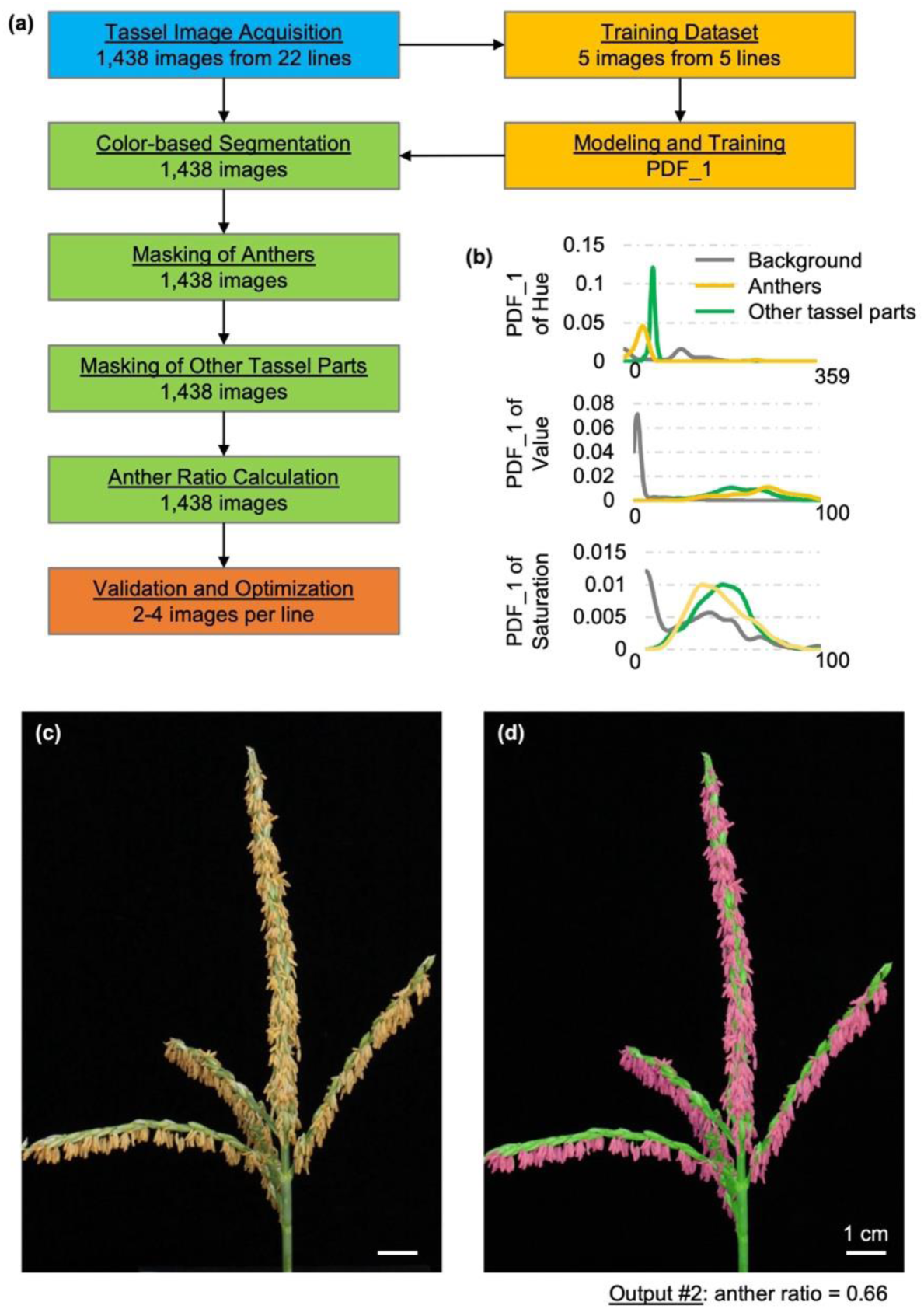
Overview of Tasselyzer. **(a)** A workflow of Tasselyzer in this study. A total of 1,438 input raw tassel images from 22 maize lines were captured. Five images were selected to generate PDF_1 as parameters for multiclass segmentation into non-plant background, anthers, and other tassel parts based on color at pixel level. PDF_1 was used to parameterize segmentation, generate masks for anthers (fuchsia) and other tassel parts (green), and calculate anther ratios for all images. **(b)** PDF_1 is a three-dimension model with statistical information of hue, saturation and value (HSV) to parameterize color-based segmentation. An example of raw tassel image of fast-flowering mini-maize (FFMM) as input in **(c)**, and pseudo-colored output image in **(d)**. Anther ratio of this example image is indicated below panel **(d)**. Bars in **(c** and **d)**, 1 cm.

Considering a tassel image from fast-flowering mini-maize (FFMM) for example, the original input image displayed a tassel with one central main spike and multiple side branches. Yellow anthers exerted from florets of all branches. After processing the image through Tasselyzer, anthers were highlighted in fuchsia and other tassel parts in bright green (Figure 1c-d). In a separate output file, Tasselyzer computed the corresponding anther ratio by summing the pixels of anthers and dividing by the total number of pixels in both anthers and other tassel parts. We propose that this automated measurement closely mirrors visual assessment made of human observers, irrespective of the size and complexity of maize tassels.

### Tasselyzer is a sensitive tool to measure anther exertion of regional or whole maize tassels

Development of a typical tassel spans 2-7 days, with the duration determined by genetic context and growth conditions. Anthers first emerge from upper florets in the center of the main spike on day 1, gradually spreading to lower florets and tips of all branches until the lowest side branch in subsequent days (Egger and Walbot, 2015). To assess if anther ratio, as determined by Tasselyzer, correlates with degree of anther exertion, we examined tassel images during anthesis. We captured and analyzed photos of developing inbred line A632 to observe the process of anthesis, and Tasselyzer demonstrated sensitivity in capturing changes in anther exertion during this period (Supporting Figure 1).

An intriguing characteristic of inbred maize lines is their highly synchronized flowering time, if environmental and artificial impacts are normalized. To further evaluate Tasselyzer across different days during anthesis, we staggered planting of 6 to 7 FFMM seeds every day over two weeks. We harvested 75 tassels and took 288 images on the same day, providing snapshots of tassels at various stages with differing degrees of anther exertion (Supporting Table 2). The width of the main spike of the tassel has traditionally been used as a proxy for tracking anther exertion (Fonseca *et al*., 2003, Mirnezami *et al*., 2021), so we manually measured main spike widths in these images using ImageJ (Schindelin *et al*., 2015; Rueden *et al*., 2017). Main spike widths of FFMM ranged from 0.38 cm to 1.45 cm. Notably, plants planted later (days 11-14), in which anthesis had not initiated, exhibited thinner main spikes, while plants planted on days 1 to 10 showed increasing and fluctuating main spike widths, consistent with anthesis initiation (Figure 2a and Supporting Figure 2). This observation aligned with findings from previous studies (Mirnezami *et al*., 2021).

**Figure 2.**
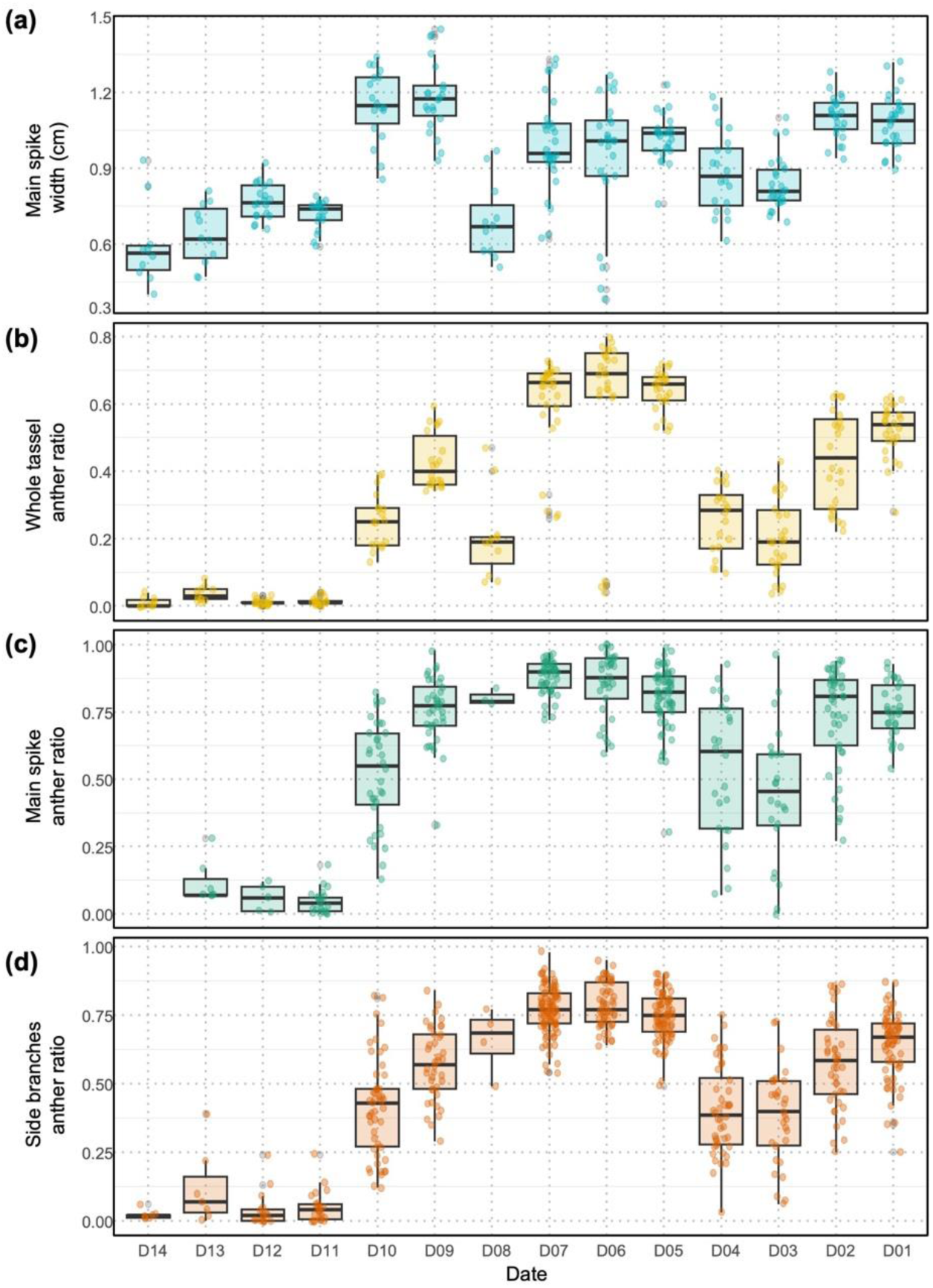
Comparison of measurements of anther exertion in FFMM on different days of anthesis. **(a)** Box and scatter plots of main spike widths were measured from tassel images with different planting dates, with D01 the earliest and D14 the latest. Box and scatter plots of anther ratios of whole tassels **(b)**, main spikes **(c)**, and side branches **(d)** were calculated using Tasselyzer. Each dot represents a value extracted from one tassel image and each box depicts a distribution of values on a planting date. All values were extracted from the same set of 288 tassel images of FFMM prepared as indicated in Materials and Methods and Supporting Table 2.

Subsequently, we applied Tasselyzer to calculate the anther ratios of whole tassels of FFMM and compared these ratios with main spike widths. Plants planted on days 11-14, which had minimal anther presence on the day of photo shooting, exhibited anther ratios close to zero, consistent with the original images (Figure 2b and Supporting Figure 3). As expected, plants planted on days 9 and 10, in which anther exertion commenced, showed anther ratios ranging from 0.18 to 0.50. Plants planted on days 5-7 exhibited medium anther ratios peaking around 0.70. The trend of anther ratio changes in FFMM is similar to that of developing tassels of A632 and A619 (Supporting Figure 1 and Supporting Figure 4). Comparing anther ratios with main spike widths, while both measurements were consistently low on days 11-14, their peaks occurred on different days. Main spike widths were less sensitive to anthesis stages, whereas anther ratios exhibited greater sensitivity to changes in anther exertion (Supporting Figure 5a). Notably, unexpectedly low values of main spike widths and anther ratios on day 8 were consistent with each other but were outliers, likely due to an error or delay in germination or growth on that day.

Furthermore, analyzing either main spike width or the entire tassel ignores regional phenotypic differences resulting from short-term developmental and environmental factors affecting anther development temporarily. To assess the performance of Tasselyzer in capturing this information, we cropped the photos of main spike and side branches from FFMM whole tassel images during anthesis and analyzed them separately with Tasselyzer (Figure 2c-d and Supporting Figure 6). For plants planted on days 11-14, anther ratios of main spikes and side branches ranged from 0 to 0.28 and 0 to 0.39, respectively. Anther ratios increased to 0.13-0.79 and 0.12-0.82 for the main spike and side branches of plants planted on day 10, right after initiation of anthesis. Both trends in regional anther ratios were comparable but more sensitive in capturing the degrees of anther exertion over anthesis, compared to the measurements of main spike widths or anther ratios of whole tassels (Supporting Figure 5b-e).

### Tasselyzer distinguishes anthers with distinct colors from other tassel parts

We then examined whether Tasselyzer could be extended to a broader selection of maize lines, using 1,438 tassel photos from 22 maize lines including 20 NAM founder lines (Supporting Table 1). To evaluate the performance of Tasselyzer, we employed standard metrics of recall, precision, and F_1_ score, a balanced score of precision and recall, through manual curation of subsets of pseudo-colored output images from each maize line (Materials and Methods and Supporting Figure 7). Our finding revealed variation in Tasselyzer’s performance across these maize lines, with scores ranging from 0.28 to 0.97. Notably, six lines — CML69, A619, FFMM, P39, Ky21, and Oh43 —exhibited high F_1_ scores, indicating robust performance (Figure 3a, Supporting Figure 8, Supporting Figure 9, and Supporting Figure 10). Notably, no P39, Ky21, and Oh43 images were used in training. These lines share a common color pattern in their anthers, facilitating accurate segmentation by Tasselyzer with PDF_1. In general, recall scores were high, around 0.9, except for Mo17, in which anthers were poorly identified due to their green color, making them difficult to distinguish from other tassel parts (Supporting Figure 9). However, precision varied across the other 16 lines, ranging from 0.16 to 0.67, owing to different and variable color patterns (Supporting Figure 10).

**Figure 3.**
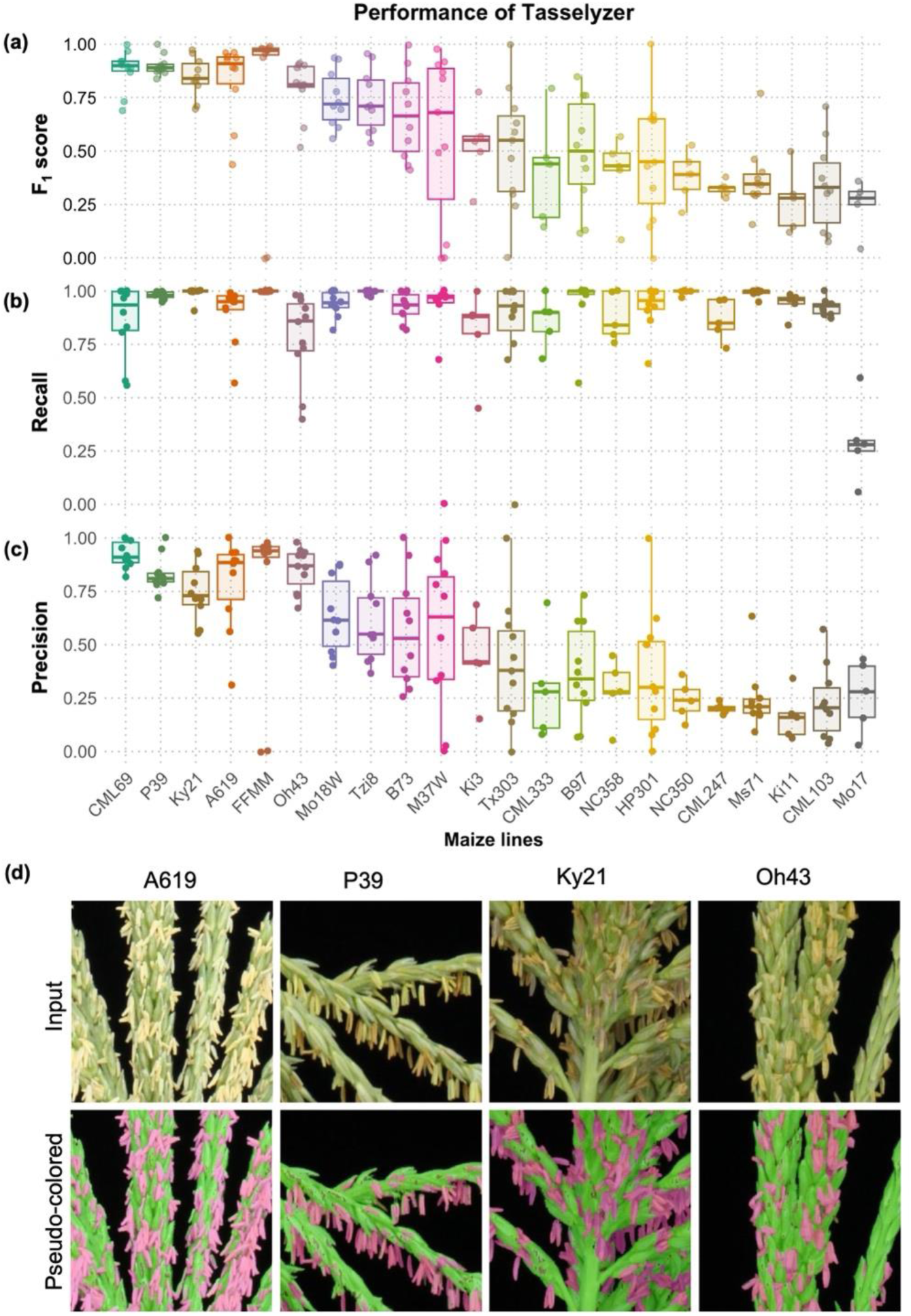
Segmentation performance of Tasselyzer on maize lines. Box and scatter plots of F_1_ score **(a)**, recall **(b)**, and precision **(c)** of Tasselyzer on individual maize line with scores ranging from high to low (left to right). Each dot represents a value extracted from one image and each box depicts a distribution of values of an individual line. **(d)** Examples of input (upper) and pseudo-colored (lower) tassel images of A619, P39, Ky21, and Oh43. More examples are in Supporting Figure 7 and Supporting Figure 8.

Assuming consistent color patterns within each maize line, a line-specific PDF (probability distribution function) could potentially enhance segmentation accuracy. To test this hypothesis, we selected one out of 30 images from NC358 and generated an NC358-specific PDF (PDF_2) for parameterizing segmentation. PDF_2 notably improved precision from 0.05-0.45 to 0.4-1.0 and F_1_ score from 0.15-0.79 to 0.46-0.93, while maintaining comparable levels of recall (Figure 4 and Supporting Figure 11).

**Figure 4.**
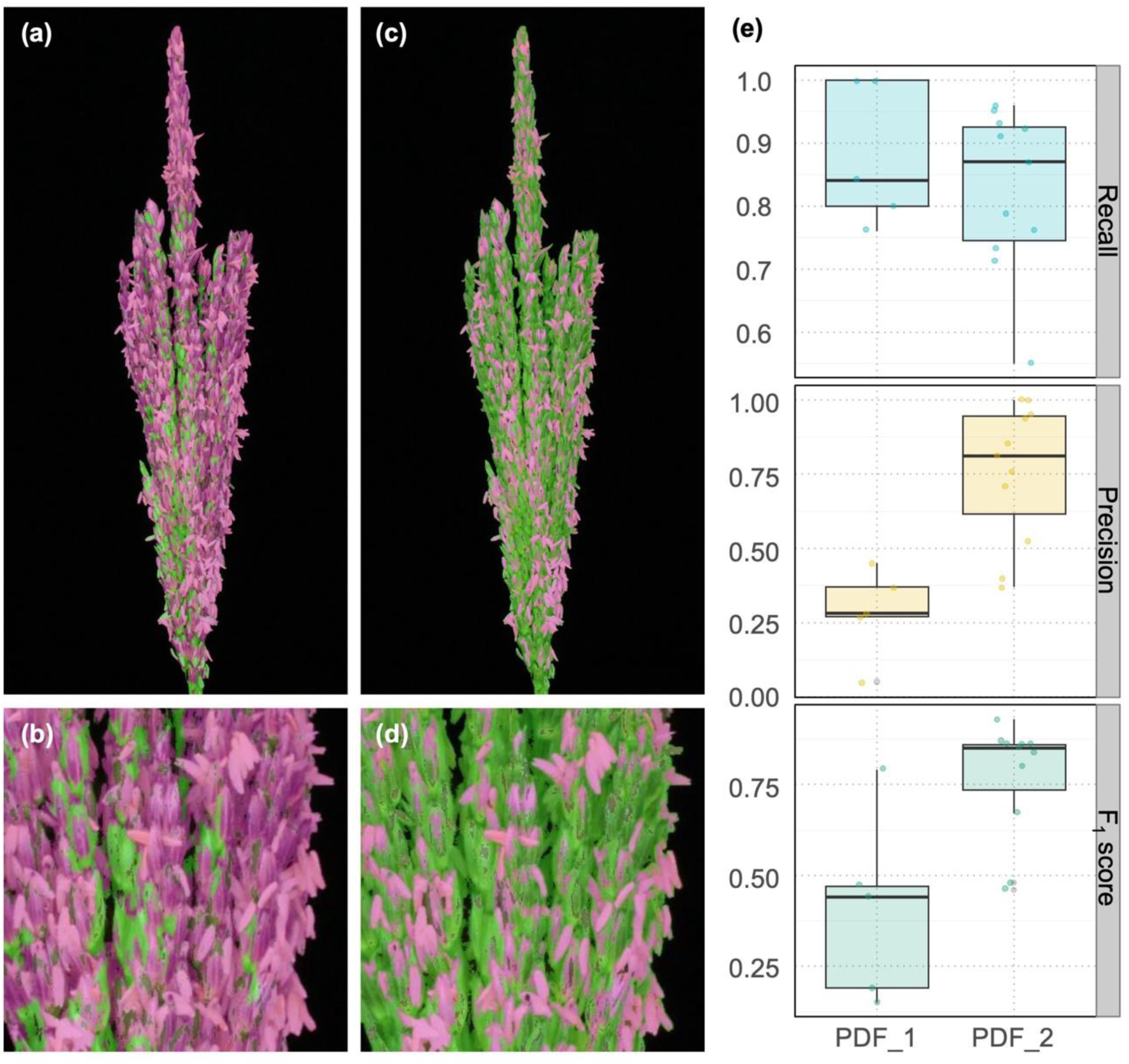
Segmentation optimization with line-specific parameter in maize line NC358. Overall and magnified images of segmentation in NC358 before **(a, b)** and after **(c, d)** optimization. **(e)** Comparison of recall, precision, and F_1_ score before (PDF_1) and after optimization with NC358 line-specific parameters (PDF_2). Scores before and after optimization with NC358-specific parameters PDF_2 were subjected to statistical analysis for significance using a one-way ANOVA test. *P*-value of difference in precision is 0.00101. *P*-value of difference in F_1_ score is 0.00377. More example images are in Supporting Figure 11.

## Discussion

The degree of anther exertion serves as a crucial indicator of male fertility in maize; however, the lack of an automated tool for quantification has presented a challenge. To address this gap, we developed an image-based system capable of recording both whole and regional tassel phenotypes. Thus, we introduced Tasselyzer, a user-friendly and cost-effective machine learning method for extracting and quantifying anther exertion information based on pixel color. This system has the capacity to process thousands of images using parameters generated from simple training steps. Like any method, Tasselyzer has advantages and limitations which users should consider when deploying it. Below, we will discuss the shortcomings and potential applications of Tasselyzer in various scenarios.

### Tasselyzer is a sensitive color-based tool to identify anthers

Tasselyzer, as a sensitive color-based tool, effectively distinguishes anthers from other parts of the tassel and a dark, non-plant background. However, its performance may vary depending on the distinctiveness of the anther color compared to other tassel parts. In our testing with 22 maize lines, Tasselyzer demonstrated acceptable performance in terms of recall, precision, and F_1_ score, with potential for improvement through optimized segmentation parameters, except for the Mo17 line. Anthers of Mo17 were green like other tassel parts, which presented a challenge to Tasselyzer; this may be the case for other lines with indistinguishable anther colors. Additionally, Tasselyzer is limited by its inability to exclude leaves or remove stems beneath the lowest branch. Currently, manually stem trimming and removal or masking of leaves before imaging is required. Theoretically, this limitation could be addressed through pre-staining anthers with specific chemicals or using specialized cameras.

Moreover, deep learning models like Segment Anything Model (SAM) potentially could address issues in distinguishing anthers (Kirillov *et al.,* 2023). Initial trials with SAM showed potential, although SAM tends to treat the entire tassel as a single object rather than segmenting out anthers (Supporting Figure 12). While this capability is useful for separating tassels from the background, it does not meet the requirement for quantifying anther exertion directly. Cropping whole tassel images into parts partially alleviates this limitation, and SAM performed similarly to the current version of Tasselyzer in terms of segmenting anthers and excluding leaf or extra stem areas (Supporting Figure 13). While SAM exhibits promise for future enhancements of Tasselyzer, its primary function is not tailored to quantifying anther exertion. Therefore, it remains outside the main scope of this study.

### Tasselyzer has potential as a non-destructive method for field studies

In this study, most of the tassel images were captured from detached tassels from either field or greenhouse grown plants and were taken in a closed photo booth to ensure uniformity of the imaging background and simplify downstream performance evaluation. To further enhance its applicability in field settings, a color card could be included during image acquisition, and additional color correction methods could be employed for color and size normalization in field-captured photos. This approach would be particularly valuable when combined with SAM or similar tools, which can segment tassel parts from the background before employing Tasselyzer to segment anthers.

### Meaning and usage of anther ratio

The size and fitness of anthers depend on genetic and environmental factors and vary among individual plants. Furthermore, old anthers drop off the tassel or senescence, and young anthers exert during a 2-7-day process of anthesis. All these factors contribute to difficulties in accurate quantification of anther exertion, unlike many other biological measurements including width, length, weight, etc. Thus, we propose recording and measuring anther exertion by taking snapshots during anthesis and calculate anther ratio. The anther ratio represents the proportion of anthers in an image compared to the entire tassel at a pixel level. Conceptually, it ranges from 0 to 1, where 0 signifies no anther exertion and 1 indicates that anthers completely cover other tassel parts. It’s important to note that the anther ratio does not provide an exact count of individual anthers; rather, it offers proportion of anthers to the whole or partial tassel for comparative purposes. For instance, it can be used to compare mutant phenotypes under different environmental conditions or to track changes during different stages of anthesis.

In our prior work and its initial application, Tasselyzer was successfully deployed to identify specific stages of maize development, such as over 9 days of meiosis and post-meiosis in maize dcl5 mutant research (Teng et al., 2020). Similarly, in this study, the anther ratio was used to monitor the anthesis process across multiple inbred lines (Figure 2). Theoretically, comparisons of anther ratios among different maize inbred lines can be made when F_1_ scores are 0.8 or higher. However, users should exercise caution when significant variations in tassel color or architecture exist, as these factors may affect the reliability of the comparison.

In summary, Tasselyzer provides a sensitive tool to analyze images of tassels at various stages of anthesis, whether from whole tassels or tassel parts, across a diverse collection of maize lines. It enables the study of environmental or stress impacts on anthesis by quantifying degrees of anthesis, despite limitations when applied to tassels with complex tassel architectures or color patterns. These limitations can be mitigated through parameter optimization and analyzing partial tassel images instead of whole tassels. Additionally, there is potential for improvement by integrating more sophisticated algorithms, such as deep learning methods, with extensive training efforts. Overall, Tasselyzer is an accessible, cost-effective, and automated tool for quantifying maize anther exertion across a diverse range of maize lines, offering valuable insights into plant development and responses to environmental factors.

## Materials and Methods

### Plant material and growing conditions

Maize (*Zea mays*) NAM founder lines (McMullen *et al*., 2009), including B73, B97, Ky21, M37W, Mo17, Ms71, Oh43, CML103, CML247, CML69, CML333, Ki3, Ki11, NC350, Tx303, Tzi8, HP301, P39, Mo18W and NC358 were cultivated at the field research site of the Donald Danforth Plant Science Center (4195 MO-94, St Charles, MO, 63301, USA) in the summer of 2023. Inbred lines including A619, A632, and FFMM (McCaw *et al*., 2016) were grown in optimized greenhouses for maize (28° C/ 22° C, 14 h day/10 h night or 16 h day/8 h night temperatures, approximately 500 μmol/m^2^/s) in the Plant Growth Facility at Donald Danforth Plant Science Center. To stagger planting for FFMM and A619 lines in the greenhouse, 6-7 seeds were planted daily over a two-week period. Anther exertion was monitored daily, and tassels of FFMM and A619 were detached and images were captured on the same day when anthers from plants planted on “day 10” started exerting from the center of main spikes. A summary table of lines and images is in Supporting Table 1 and Supporting Table 2.

### Tassel imaging

We implemented an imaging system designed to capture RGB images of detached tassels with multiple options of camera options and conditions, aiming to optimize image quality while minimizing cost, time, and computing power investment. Our final setup featured an enclosed box illuminated by LED lights (Neewer 32-inch shooting tent, Amazon), with black fabric serving as background. A DSLR camera (Canon EOS Rebel T1i) equipped with an EF-S 18-55 mm f/3.5-5.6 IS lens was employed, controlled by ggphoto2 software installed on a Raspberry Pi computer. This setup ensured consistent imaging settings, including manual mode with ISO at 800, aperture at 16, shutter speed at 1/80^th^ of a second, and resolution at 15.1-megapixel, facilitating downstream analysis and providing descriptive image names. For potential color calibration and size measurement purposes, a color card with ruler (X-Rite ColorChecker classic mini) was included in all tassel images. Each tassel was photographed at least four times, capturing variation from approximately 90-degree rotations to provide comprehensive coverage from different angles.

### Measurements of tassel main spike width

The measurement of tassel main spike width was conducted using a ruler reference provided by the X-Rite ColorChecker classic mini card in each image. The measurement was achieved by employing “straight line” and “measure” tools available in ImageJ (Schindelin *et al*., 2015; Rueden *et al*., 2017). Main spike widths of each group of FFMM plants from different planting days were subjected to statistical analysis for significance using a one-way ANOVA test, followed by *post-hoc t*-tests with Holm corrections.

### Tassel segmentation by machine learning

The Tasselyzer pipeline was implemented using naïve Bayes classifier module from the open-source software package PlantCV (Fahlgren *et al*, 2015; Abbasi and Fahlgren, 2016; Gehan *et al*., 2017; Schuhl *et al*., 2022). It was assumed that colors of the multiclass in tassel images (background, anthers, and other tassel parts) are distinct. RGB values for each class were sampled using the pixel inspection tool of ImageJ (Schindelin *et al*., 2015; Rueden *et al*., 2017) to extract color features and statistical information for generating the probability distribution functions (PDF) for training. The Maximum likelihood ratio test was then employed as the naïve Bayes approach to identify pixels, using corresponding PDF. In the initial phase of this study, 4,294 pixels were sampled for each class from five training images to generate PDF_1 (see training image information in Supporting Table 1), which was subsequently applied to segment all tassel images in this study, unless otherwise specified. To enhance segmentation performance in the NC358 line, 76 pixels were sampled for each class from one training image to generate PDF_2, which was then applied to segment the remaining NC358 images.

### Anther ratio calculation

Tasselyzer generates masks for anther and other tassel parts based on the segmentation results. These masks are then overlaid onto the original input images, with anthers depicted in pseudo-colored fuchsia and other tassel parts in green. This visualization serves for visual inspection and demonstration purposes. To calculate anther ratio, Tasselyzer sums up the total number of pixels corresponding to anther (n1) and other tassel parts (n2). The anther ratio is then computed by dividing the pixel count of anthers by the total number pixels in the tassel [Anther ratio = n1/(n1+n2)]. The anther ratios of each group of plants were subjected to statistical analysis for significance using a one-way ANOVA test, followed by *post-hoc t*-tests with Holm corrections.

### Performance evaluation of Tasselyzer

We employed standard measures including recall, precision, and F_1_ score. This assessment involved randomly selecting 2-4 pseudo-colored tassel images from results of the testing dataset. These images were cropped into smaller segments (320 x 320 pixels), focusing on main spikes, side branches, and branch zones. Anther pixels were manually labeled into regions of interests (ROIs) as true positive (TP) within anther regions with fuchsia mask, false positive (FP) of other tassel regions with fuchsia mask, false negative (FN) of anther regions with green mask using the “freehand selection” and “measure” tools of ImageJ (Schindelin *et al*., 2015; Rueden *et al*., 2017). Yellow outlines were used to highlight all ROIs (Supporting Figure 7). Subsequently, we annotated, measured, and summed the pixel numbers to calculate recall, precision, and F_1_ scores. Recall represents the fraction of correctly identified anther areas of the actual anther areas [Recall = TP/(TP+FN)], indicating the ability of Tasselyzer to locate anthers in random tassel images. Precision denotes the fraction of correctly identified anther areas among all Tasselyzer identified anther areas [Precision = TP/(TP+FP)], indicating the ability of Tasselyzer to identify anthers accurately without including other tassel parts. The F_1_ score, a balanced metric, combines recall and precision to evaluate the overall performance of Tasselyzer [F_1_ score = 2*TP/(2*TP+FP+FN)]. These scores range between 0 to 1, where scores above 0.9 are considered excellent, scores between 0.8 and 0.9 are good, scores between 0.5 to 0.8 are considered average, and scores below 0.5 indicate poor performance. The scores of each group of plants were subjected to statistical analysis for significance using a one-way ANOVA test, followed by *post-hoc t*-tests with Holm corrections.

## Code and Data Availability

Tasselyzer is based on PlantCV and developed by integrating Matplotlib v 3.2.1, PlantCV v4, OpenCV v 3.4.9, NumPy v 1.18.1, Python v 3.7.7, and skimage v 0.14.3 modules. The code, the original and pseudo-colored images in this study are available in an open-access folder with the title of this publication within the GitHub of https://github.com/danforthcenter/plantcv-tasselyzer-tutorial; also within Zenodo at https://doi.org/10.5281/zenodo.5524971 (Teng *et al*., 2021). The full image sets were used in this study are available within Zenodo at https://doi.org/10.5281/zenodo.5525073 (Teng *et al*., 2021).

## Acknowledgements

We are grateful to Gus Vogt, Nicole Burkett, the field research site staff, and the plant growth facility staff at the Donald Danforth Plant Science Center for their invaluable assistance in facilitating the growth of maize. Additionally, we would like to express our appreciation to Dr. Arash Abbasi, Haley Schuhl, Dhiraj Srivastava and Josh Rothhaupt for their assistance with coding and data handling. Special thanks to Alexander Liu and Dr. Chris Topp at the Donald Danforth Plant Science Center for providing tassels of NAM founder lines in the field. We also acknowledge Rhea Kaw, Josh Sumner and Jeffery Berry for their assistance in setting up the photo shooting system. Furthermore, we are grateful to Professor Virginia Walbot for her insightful feedback and Dr. Joanna Friesner for assisting with manuscript editing. This work was made possible by the support of the U.S. National Science Foundation Plant Genome Research Program (NSF-PGRP) awards 1649424 and 1754097, as well as the Molecular and Cellular Biosciences (NSF-MCB) award 2320971.

## Author contributions

C.T., N.F., and B.C.M. conceived of the project. N.F. programmed the scripts, while C.T. performed and interpreted experiments. C.T. authored the manuscript, with editing by N.F. and B.C.M.

## Supporting Dataset

**Supporting Figure 1.**
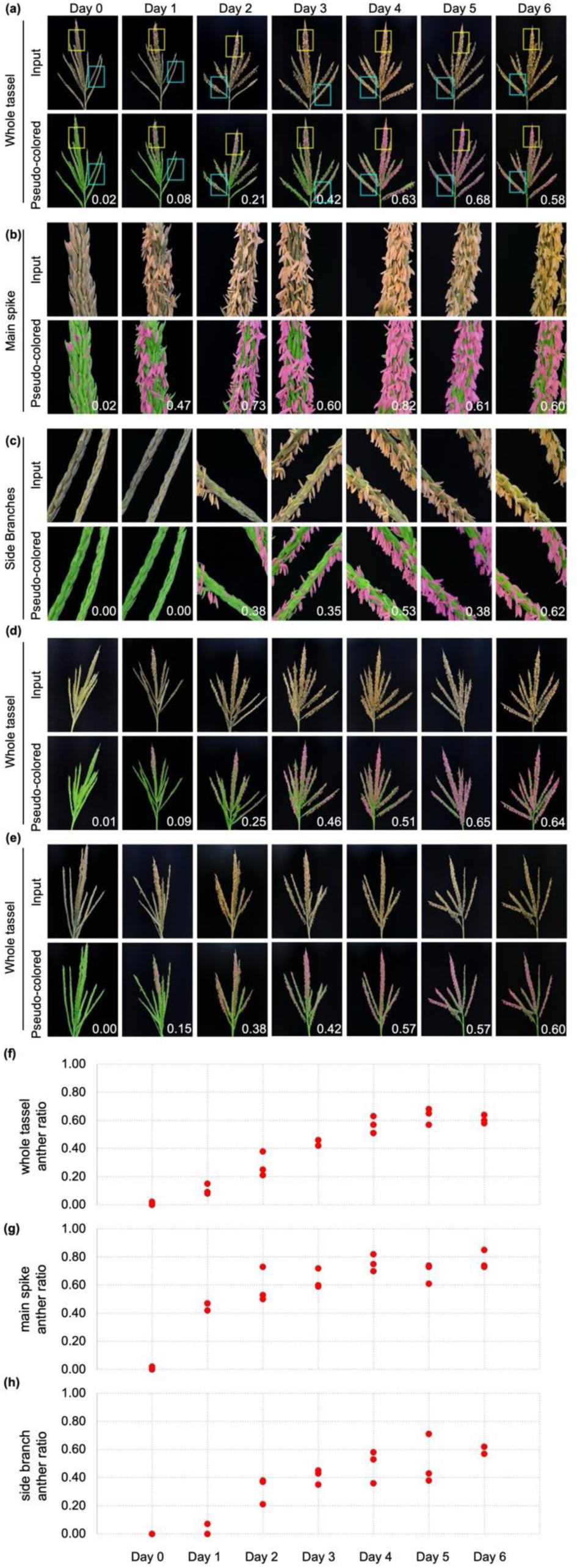
Segmentation and anther ratios during anthesis in A632 line. **(a)** Input (upper) and pseudo-colored (lower) images from an example fertile A632 inbred tassel from days 0 to day 6. Pseudo-colored images with pixels classified as other tassel parts (green) and anthers (fuchsia) overlay on the images. **(b)** Enlarged input (upper) and pseudo-colored (lower) images of main spike in (**a**, yellow boxes). **(c)** Enlarged input (upper) and pseudo-colored (lower) images of side branches in (**a**, cyan boxes). **(d, e)** Biological replicates of A632 tassels over anthesis, with input images in the upper panels and pseudo-colored images in the lower panels. Anther ratio of individual image is indicated in the bottom right and plotted in **(f)** for whole tassel, (g) for main spikes, and **(h)** for side branches.

**Supporting Figure 2.**
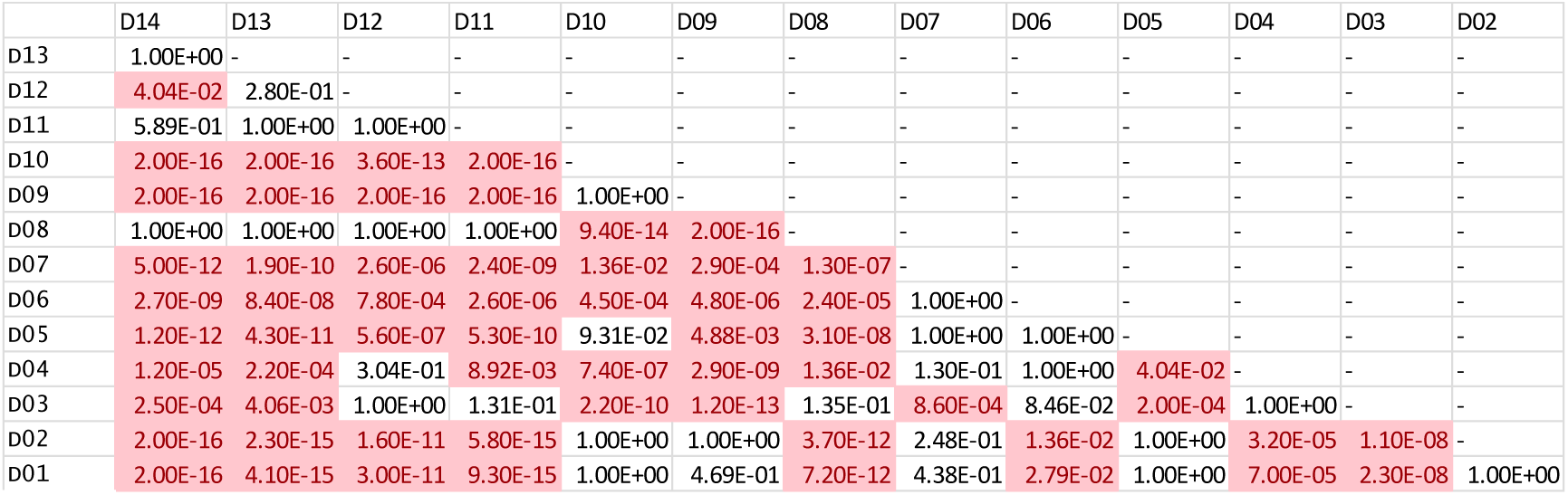
Pairwise comparison of main spike widths among groups of plants. The main spike widths of each group of plants from different planting days were tested for significance by one-way ANOVA and *post-hoc t*-test with Holm corrections. The pairs (horizontal -vertical) with significant differences (*p* < 0.05) highlighted in red.

**Supporting Figure 3.**
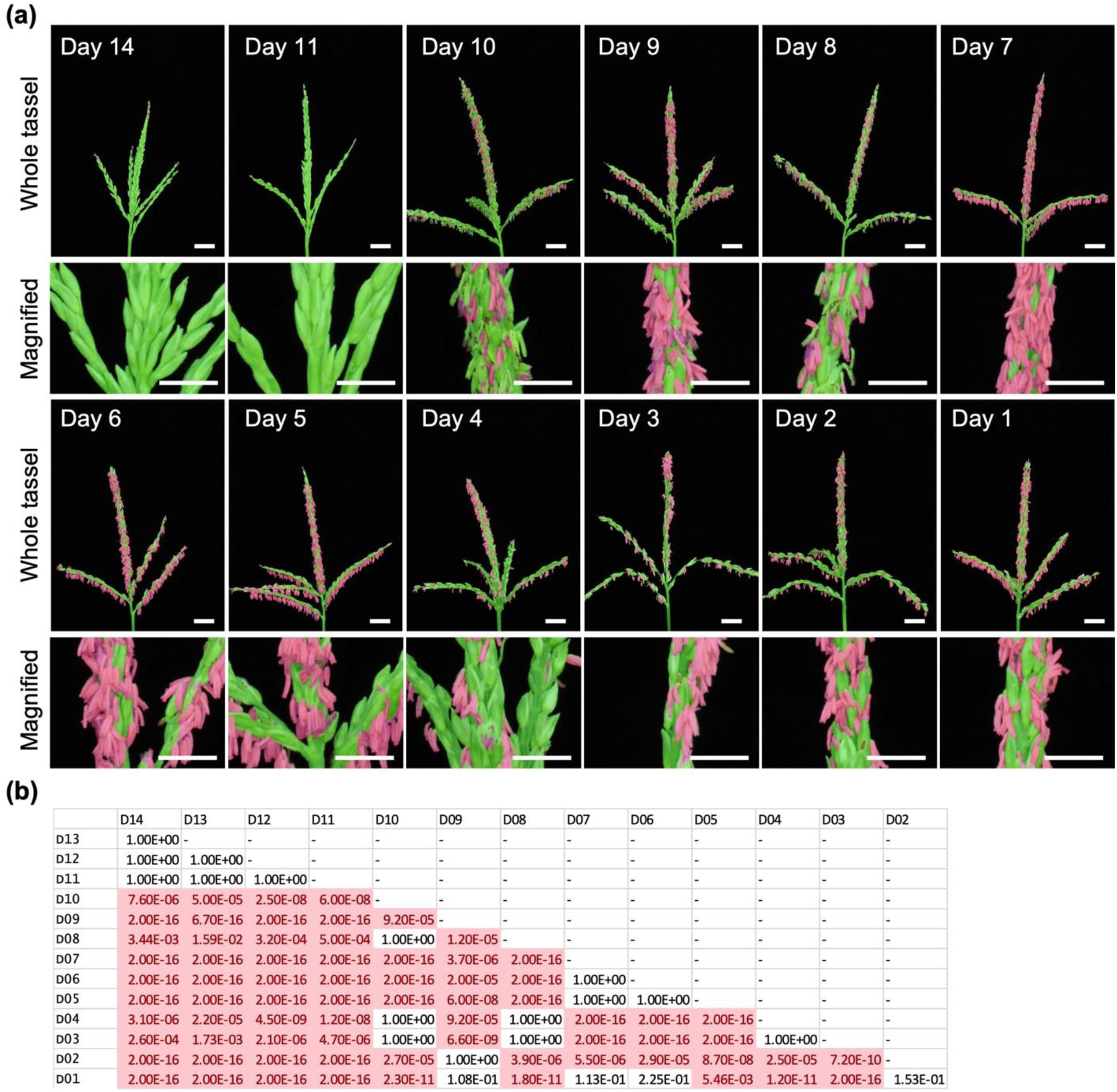
Segmentation of anthers during anthesis in FFMM line and pairwise comparison of whole tassel anther ratios among groups of plants. **(a)** Examples of pseudo-colored whole tassel (upper) and magnified (lower) images from different planting days in FFMM line. **(b)** Pairwise comparison of whole tassel anther ratios from different planting dates. The anther ratios of each group of plants from different planting days were tested for significance by one-way ANOVA and *post-hoc t*-test with Holm corrections. The pairs (horizontal -vertical) with significant differences (*p* < 0.05) highlighted in red.

**Supporting Figure 4.**
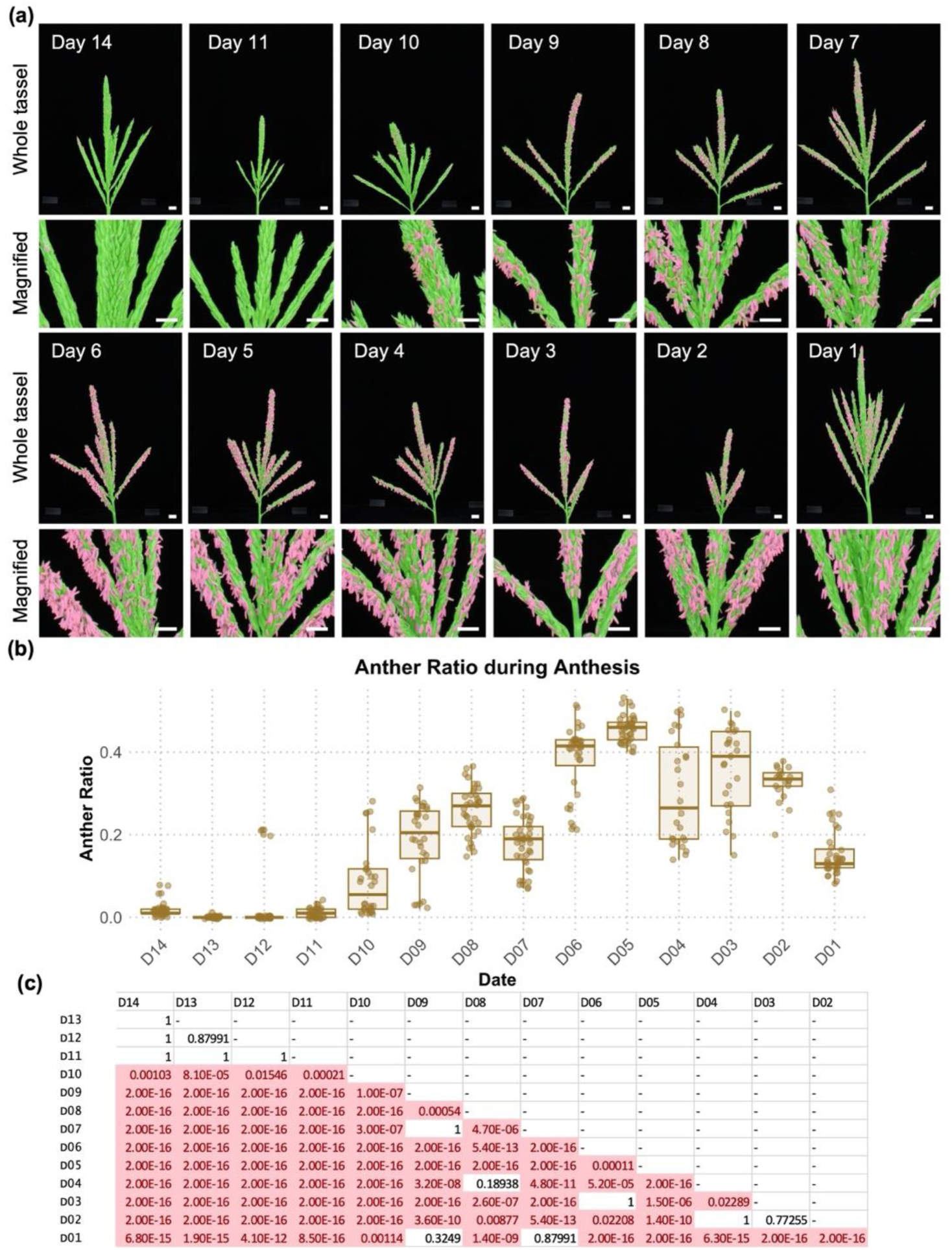
Segmentation of anthers during anthesis in A619 line. **(a)** Examples of pseudo-colored whole tassel (upper) and magnified (lower) images from different planting days in A619 line. **(b)** Anther ratios of whole tassels were calculated using Tasselyzer. **(c)** Pairwise comparison of whole tassel anther ratios from different planting dates. The anther ratios of each group of plants from different planting days were tested for significance by one-way ANOVA and *post-hoc t*-test with Holm corrections. The pairs (horizontal - vertical) with significant differences (*p* < 0.05) highlighted in red.

**Supporting Figure 5.**
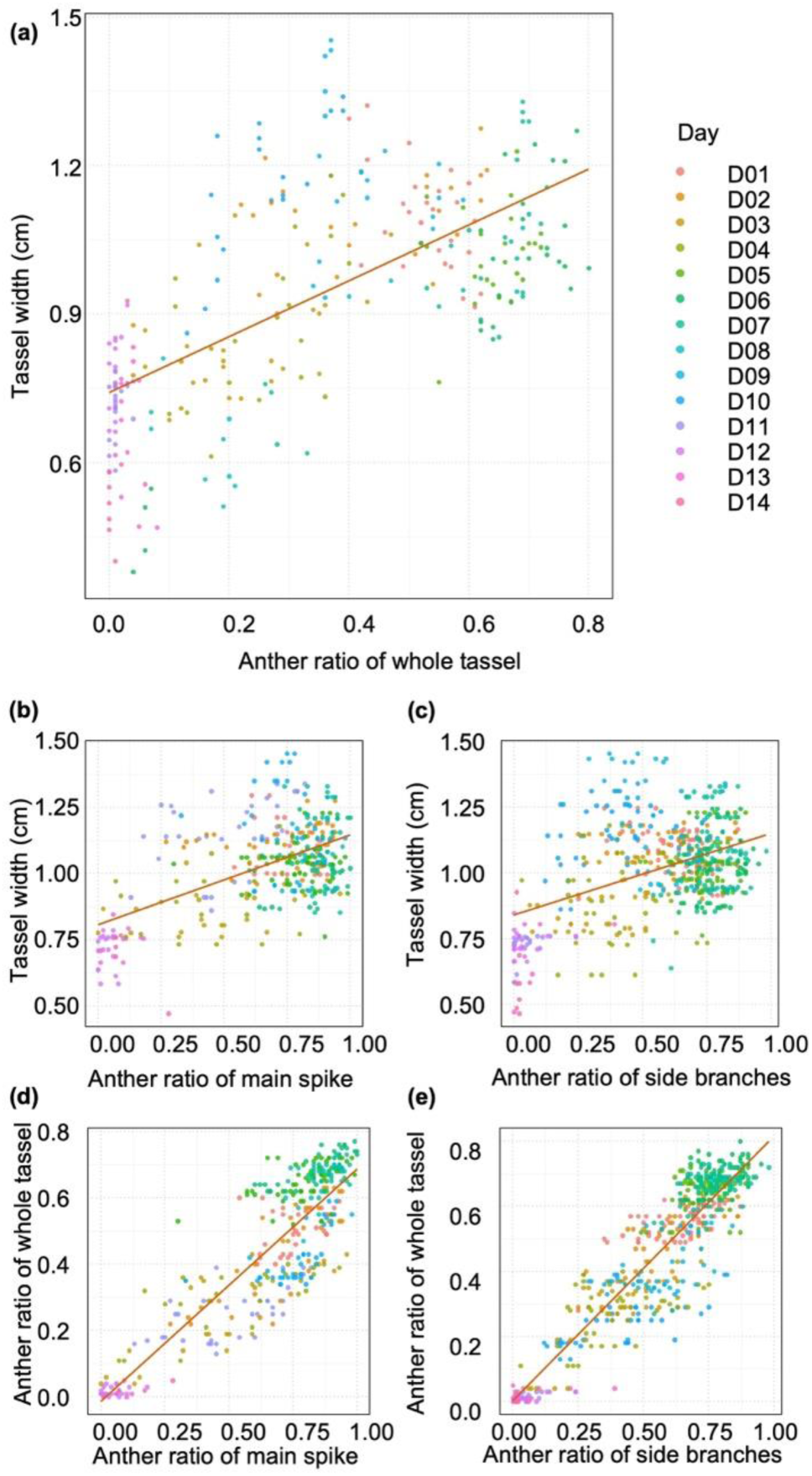
Comparison and linear regression of anther exertion measurements. **(a)** Comparison of anther ratios (x-axis) and main spike widths (y-axis) of tassels on different planting dates and linear regression (red line). Data points collected from different planting dates are in distinct colors as indicated in the legend. **(b)** Comparison of anther ratios of main spike (x-axis) and main spike width (y-axis). **(c)** Comparison of anther ratios of side branches (x-axis) and main spike width (y-axis). **(d)** Comparison of anther ratios of main spike (x-axis) and those of whole tassels (y-axis). **(e)** Comparison of anther ratios of side branches (x-axis) and those of whole tassels (y-axis).

**Supporting Figure 6.**
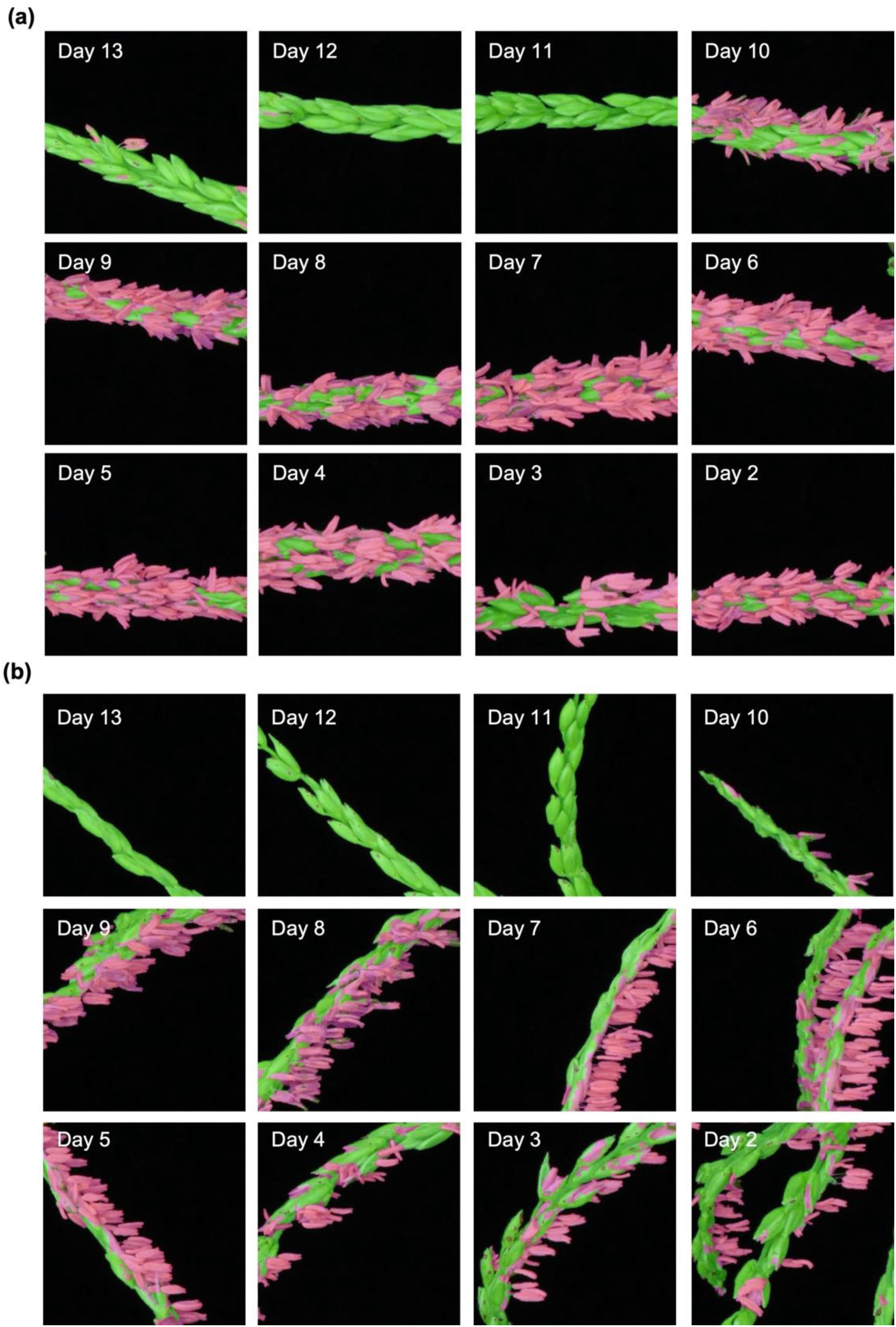
Segmentation of anthers in the tassel parts during anthesis in FFMM line. **(a)** Examples of pseudo-colored main spike images. (b) Examples of pseudo-colored side branch images.

**Supporting Figure 7.** Regions of interest (ROI) and sums in processed tassel images for performance evaluation with PDF_1 for individual maize lines. **(a)** FFMM, **(b)** A619, **(c)** Mo18W, **(d)** CML619, **(e)** M37W, **(f)** B73, **(g)** P39, **(h)** Ky21, **(i)** Tzi8, **(j)** Oh43, **(k)** HP301, **(l)** Tx303, **(m)** Ms71, **(n)** B97, **(o)** CML103, **(p)** NC358), **(q)** Ki11, **(r)** CML247, **(s)** NC350, **(t)** CML333, **(u)** Mo17, **(v)** Ki3. True positive (TP) of anther regions with fuchsia mask, false positive (FP) of other tassel regions with fuchsia mask, false negative (FN) of anther regions with green mask were manually labeled, annotated, and calculated using ImageJ (Schindelin *et al*., 2015; Rueden *et al*., 2017). Sum of TP, FP, FN, and calculated recall, precision, and F_1_ scores are listed in Supporting Dataset. Due to the limit of displaying panels per page, individual panel and legend are listed in the following pages.

**Supporting Figure 7a.**
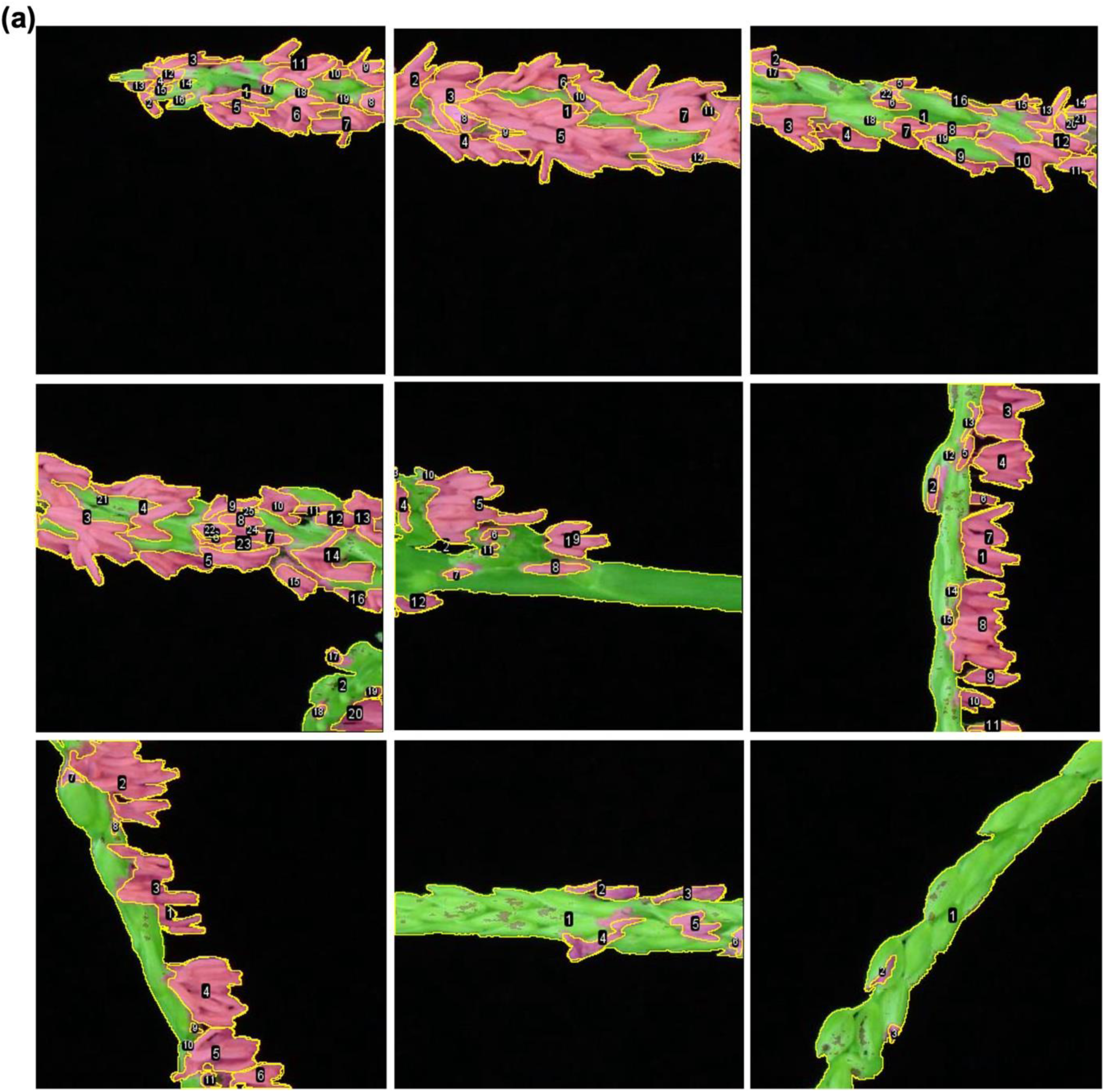
ROI and sums in processed tassel images for performance evaluation with PDF_1 for maize inbred line FFMM. TP of anther regions with fuchsia mask, FP of other tassel regions with fuchsia mask, FN of anther regions with green mask were manually labeled, annotated, and calculated with ImageJ. Sum of TP, FP, FN, and calculated recall, precision, and F_1_ scores are listed in Supporting Dataset.

**Supporting Figure 7b.**
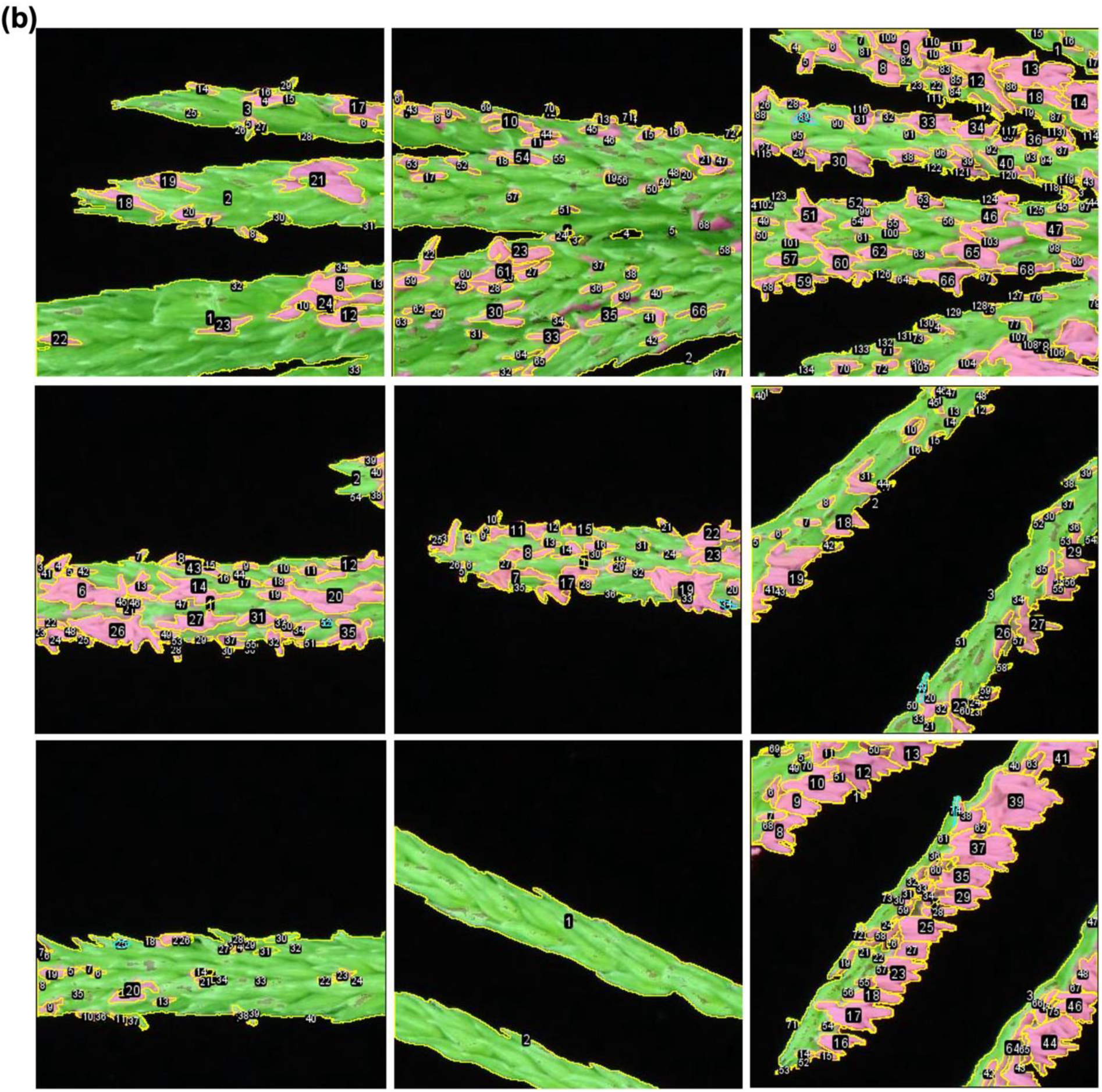
ROI and sums in processed tassel images for performance evaluation with PDF_1 for maize inbred line A619. TP of anther regions with fuchsia mask, FP of other tassel regions with fuchsia mask, FN of anther regions with green mask were manually labeled, annotated, and calculated with ImageJ. Sum of TP, FP, FN, and calculated recall, precision, and F_1_ scores are listed in Supporting Dataset.

**Supporting Figure 7c.**
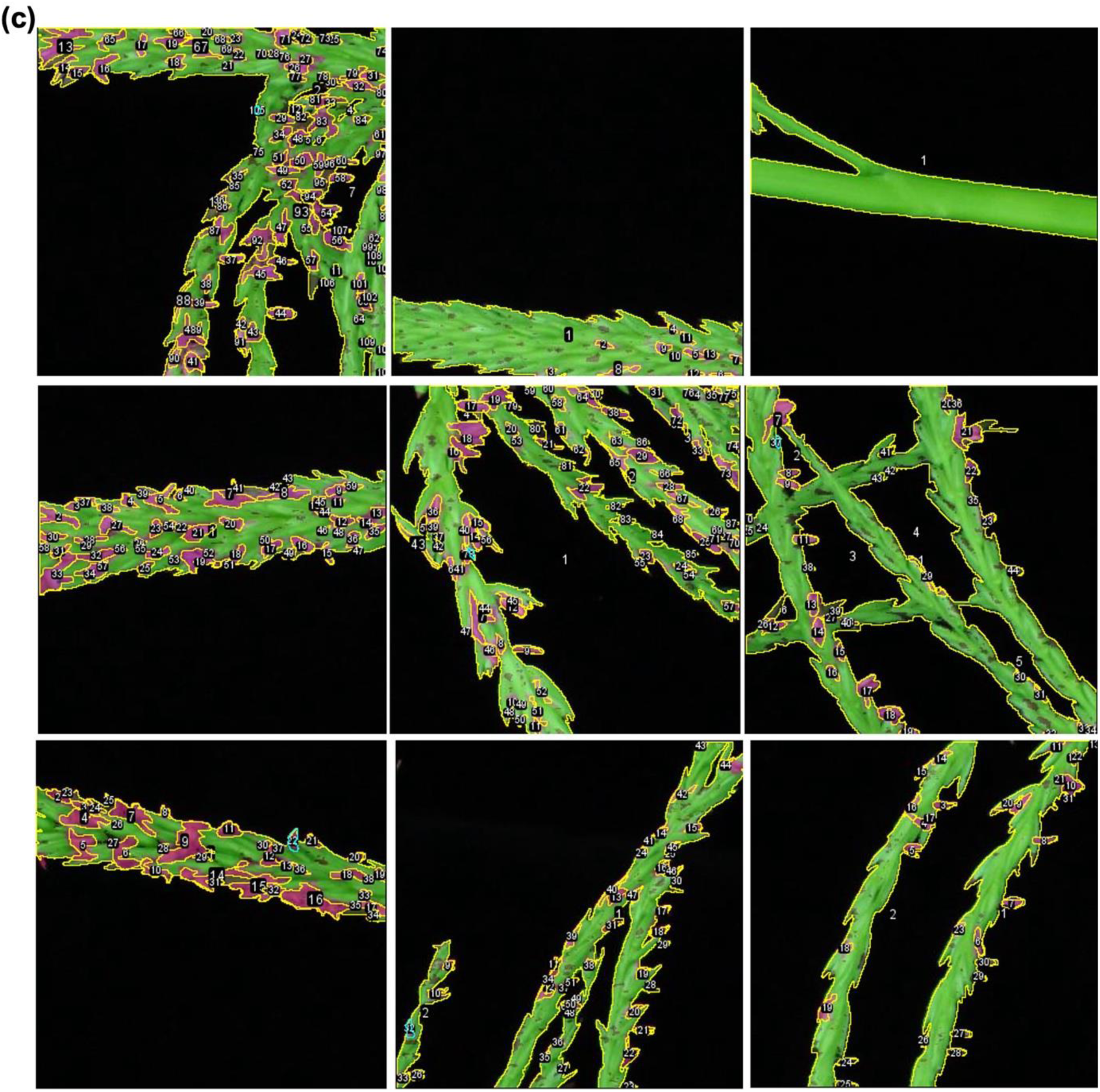
ROI and sums in processed tassel images for performance evaluation with PDF_1 for maize NAM founder line Mo18W. TP of anther regions with fuchsia mask, FP of other tassel regions with fuchsia mask, FN of anther regions with green mask were manually labeled, annotated, and calculated with ImageJ. Sum of TP, FP, FN, and calculated recall, precision, and F_1_ scores are listed in Supporting Dataset.

**Supporting Figure 7d.**
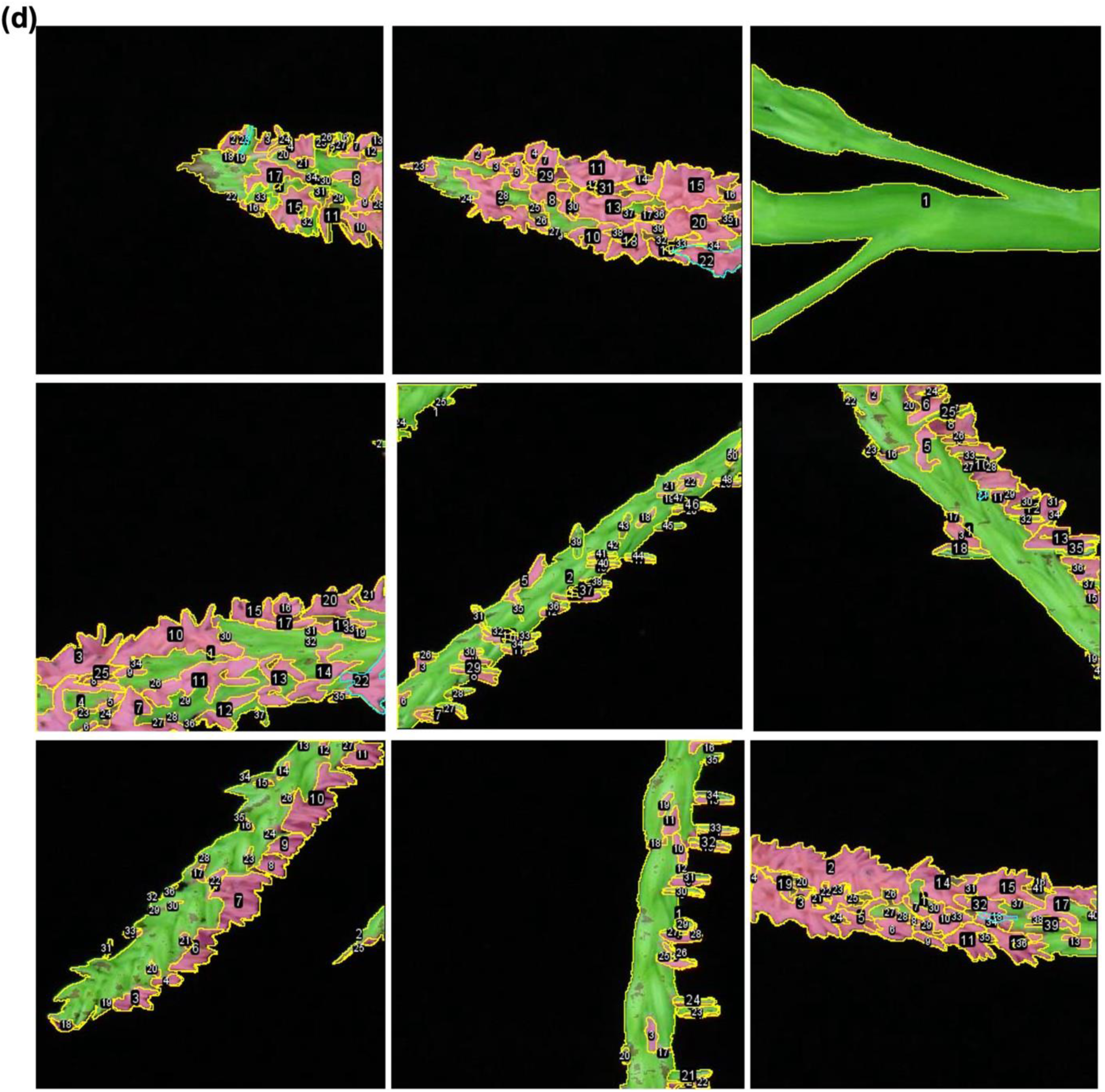
ROI and sums in processed tassel images for performance evaluation with PDF_1 for maize NAM founder line CML69. TP of anther regions with fuchsia mask, FP of other tassel regions with fuchsia mask, FN of anther regions with green mask were manually labeled, annotated, and calculated with ImageJ. Sum of TP, FP, FN, and calculated recall, precision, and F1 scores are listed in Supporting Dataset.

**Supporting Figure 7e.**
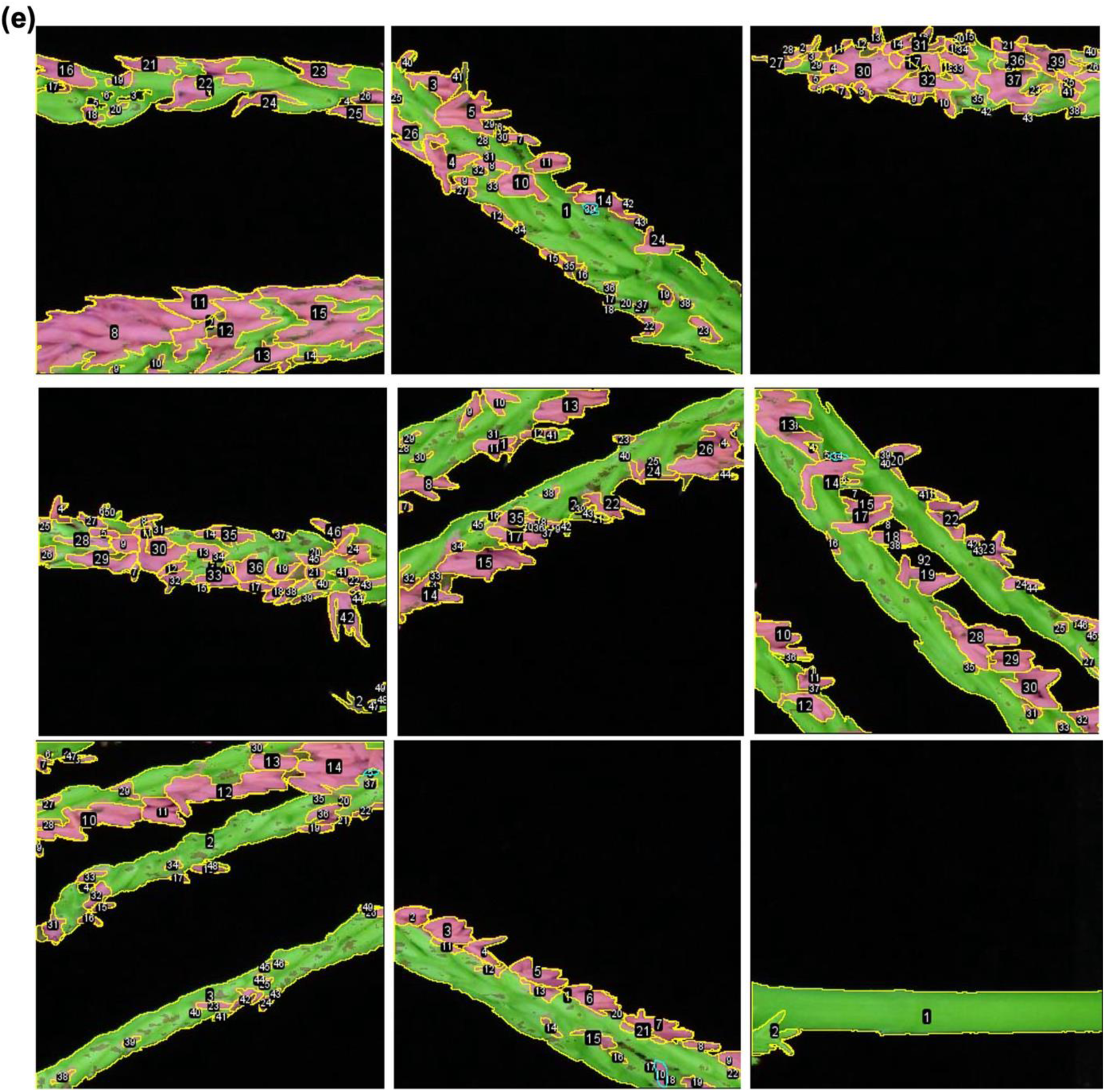
ROI and sums in processed tassel images for performance evaluation with PDF_1 for maize NAM founder line M37W. TP of anther regions with fuchsia mask, FP of other tassel regions with fuchsia mask, FN of anther regions with green mask were manually labeled, annotated, and calculated with ImageJ. Sum of TP, FP, FN, and calculated recall, precision, and F_1_ scores are listed in Supporting Dataset.

**Supporting Figure 7f.**
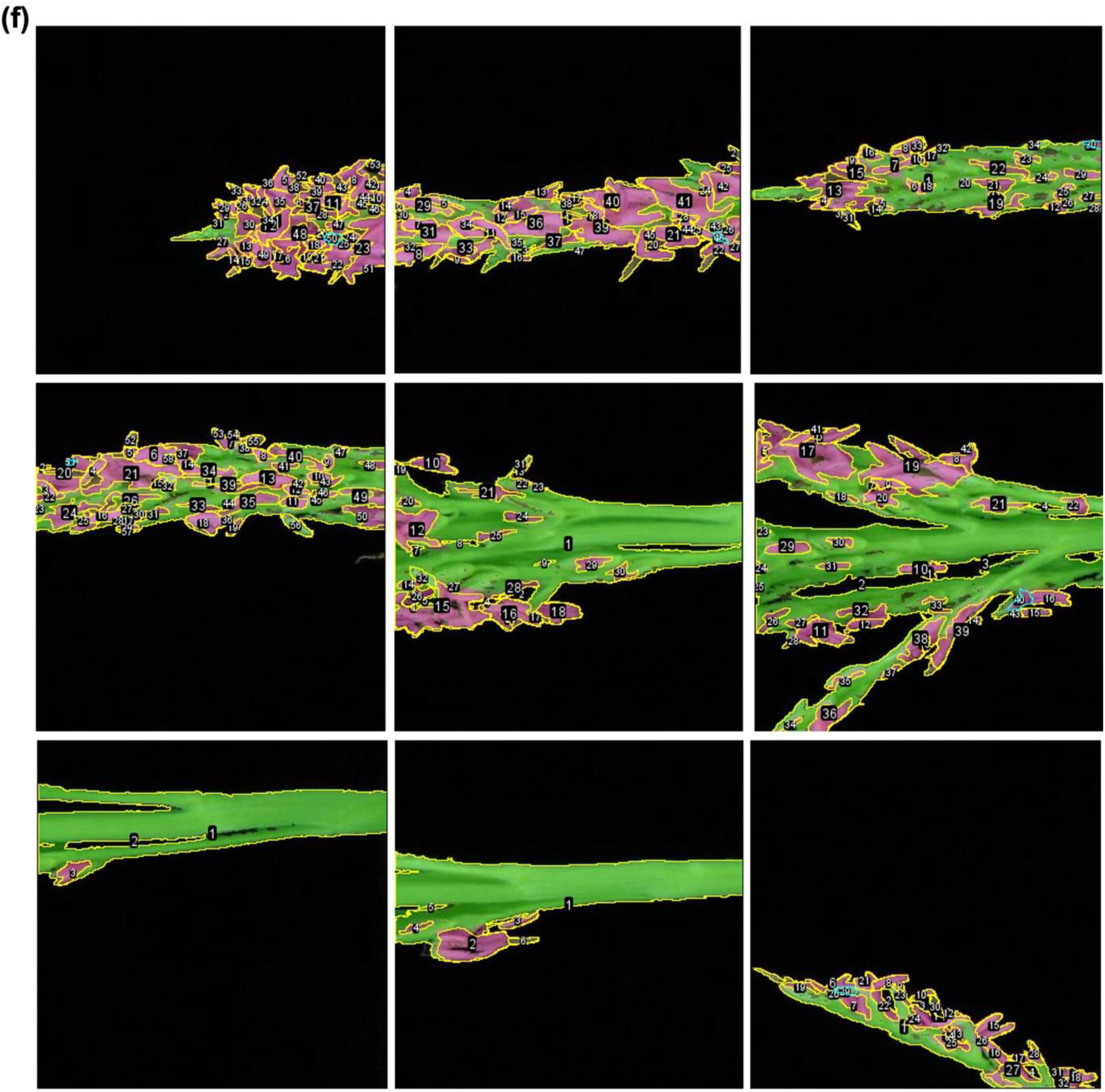
ROI and sums in processed tassel images for performance evaluation with PDF_1 for maize NAM founder line B73. TP of anther regions with fuchsia mask, FP of other tassel regions with fuchsia mask, FN of anther regions with green mask were manually labeled, annotated, and calculated with ImageJ. Sum of TP, FP, FN, and calculated recall, precision, and F_1_ scores are listed in Supporting Dataset.

**Supporting Figure 7g.**
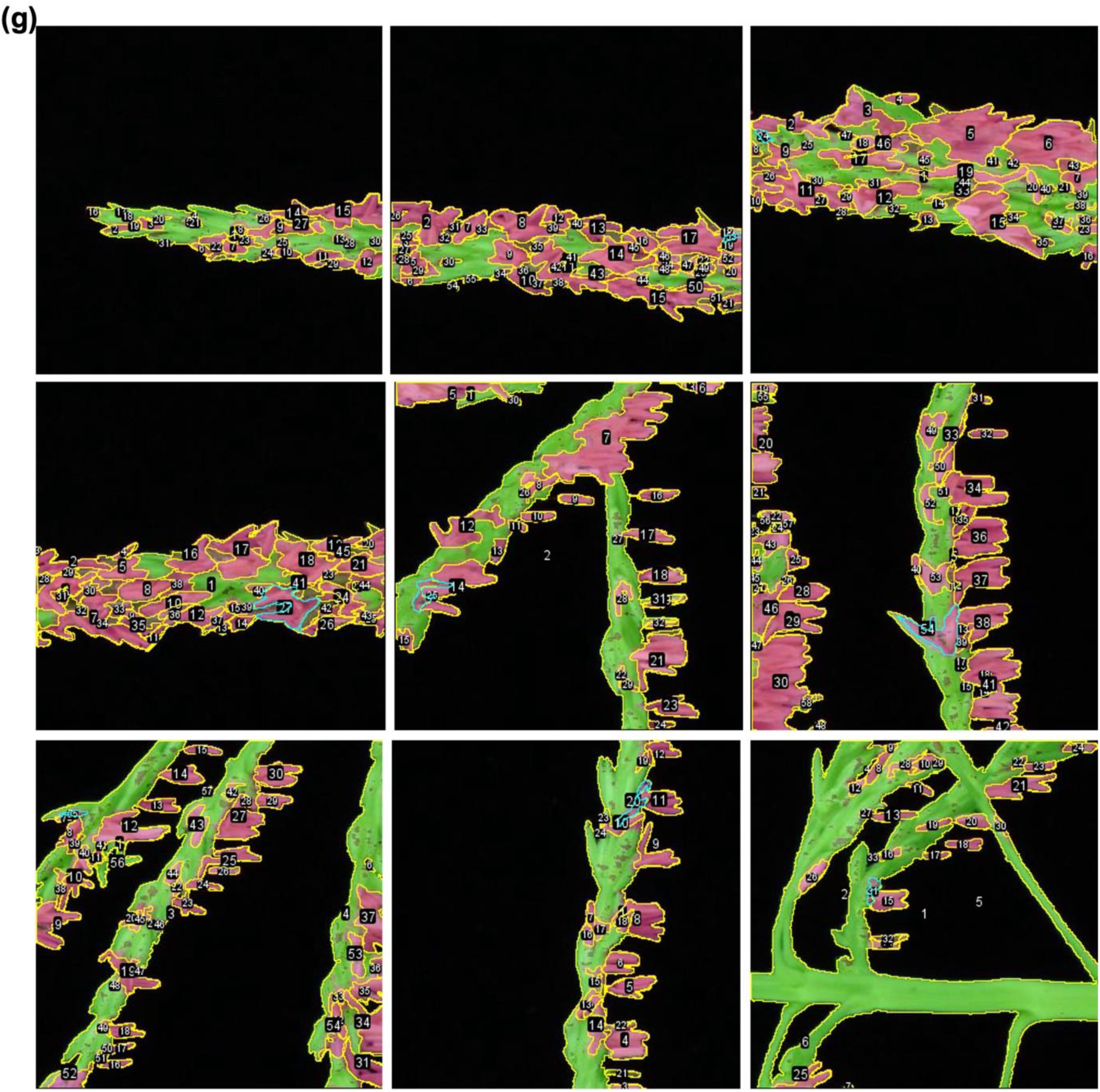
ROI and sums in processed tassel images for performance evaluation with PDF_1 for maize NAM founder line P39. TP of anther regions with fuchsia mask, FP of other tassel regions with fuchsia mask, FN of anther regions with green mask were manually labeled, annotated, and calculated with ImageJ. Sum of TP, FP, FN, and calculated recall, precision, and F1 scores are listed in Supporting Dataset.

**Supporting Figure 7h.**
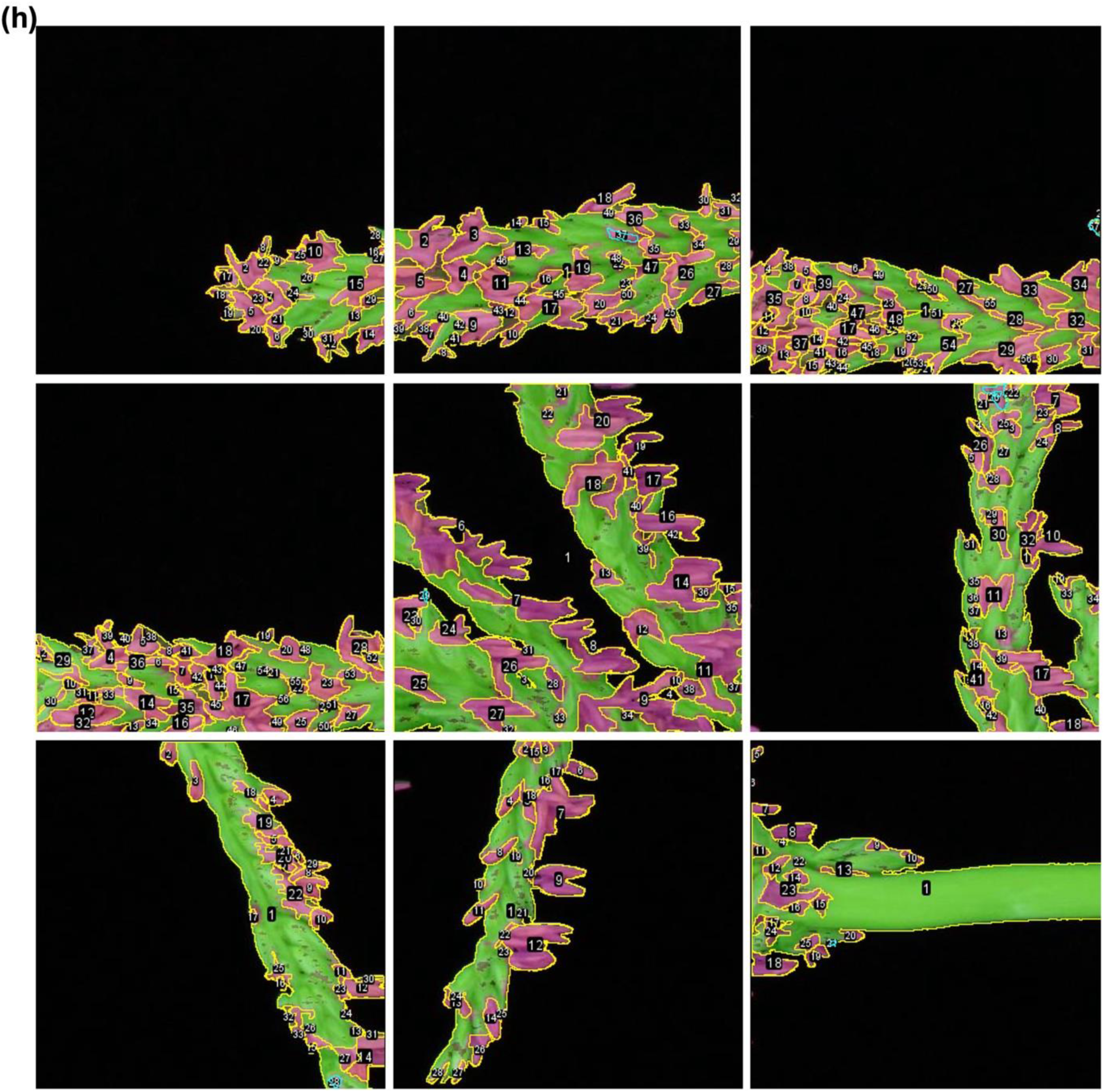
ROI and sums in processed tassel images for performance evaluation with PDF_1 for maize NAM founder line Ky21. TP of anther regions with fuchsia mask, FP of other tassel regions with fuchsia mask, FN of anther regions with green mask were manually labeled, annotated, and calculated with ImageJ. Sum of TP, FP, FN, and calculated recall, precision, and F_1_ scores are listed in Supporting Dataset.

**Supporting Figure 7i.**
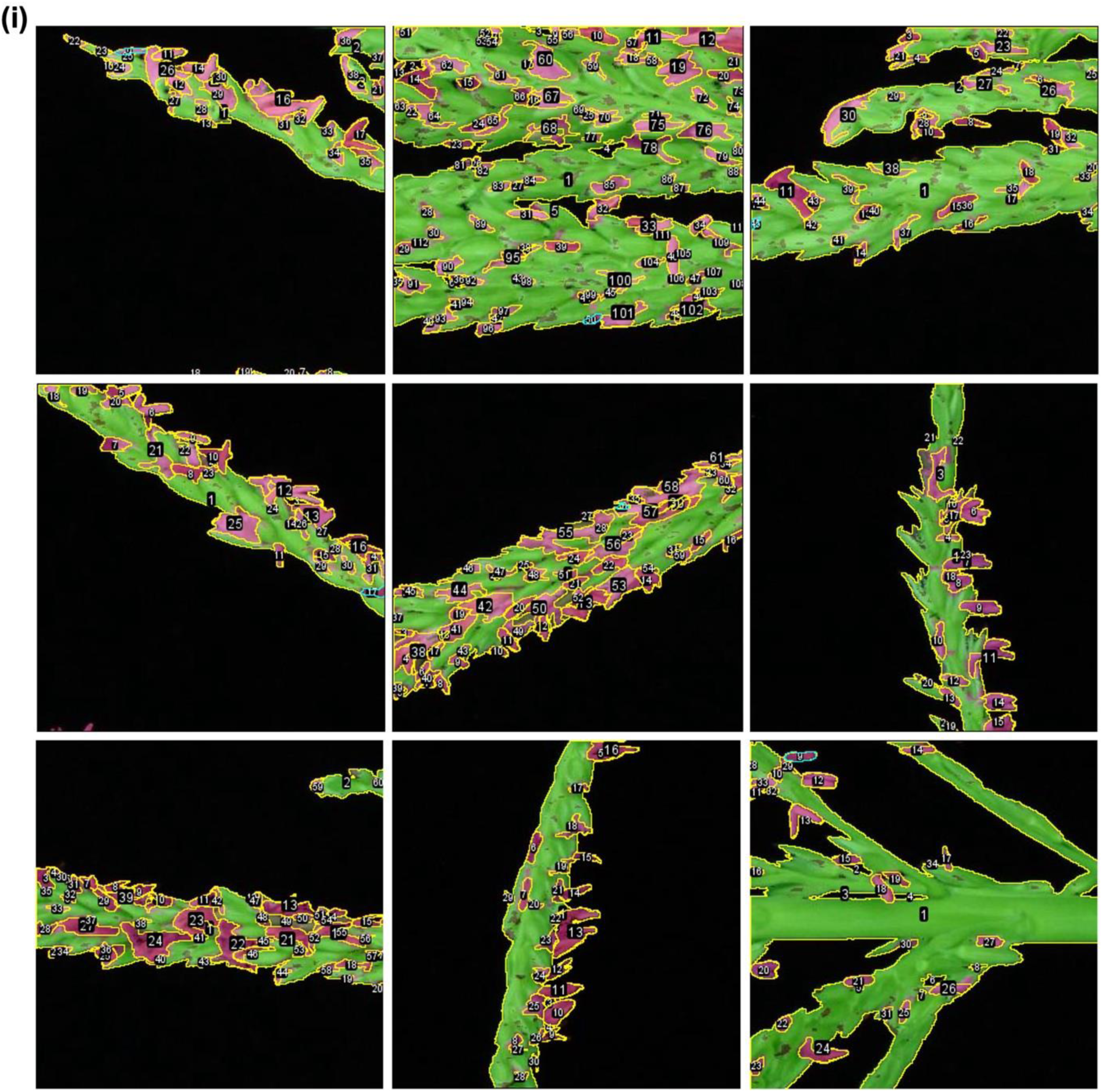
ROI and sums in processed tassel images for performance evaluation with PDF_1 for maize NAM founder line Tzi8. TP of anther regions with fuchsia mask, FP of other tassel regions with fuchsia mask, FN of anther regions with green mask were manually labeled, annotated, and calculated with ImageJ. Sum of TP, FP, FN, and calculated recall, precision, and F_1_ scores are listed in Supporting Dataset.

**Supporting Figure 7j.**
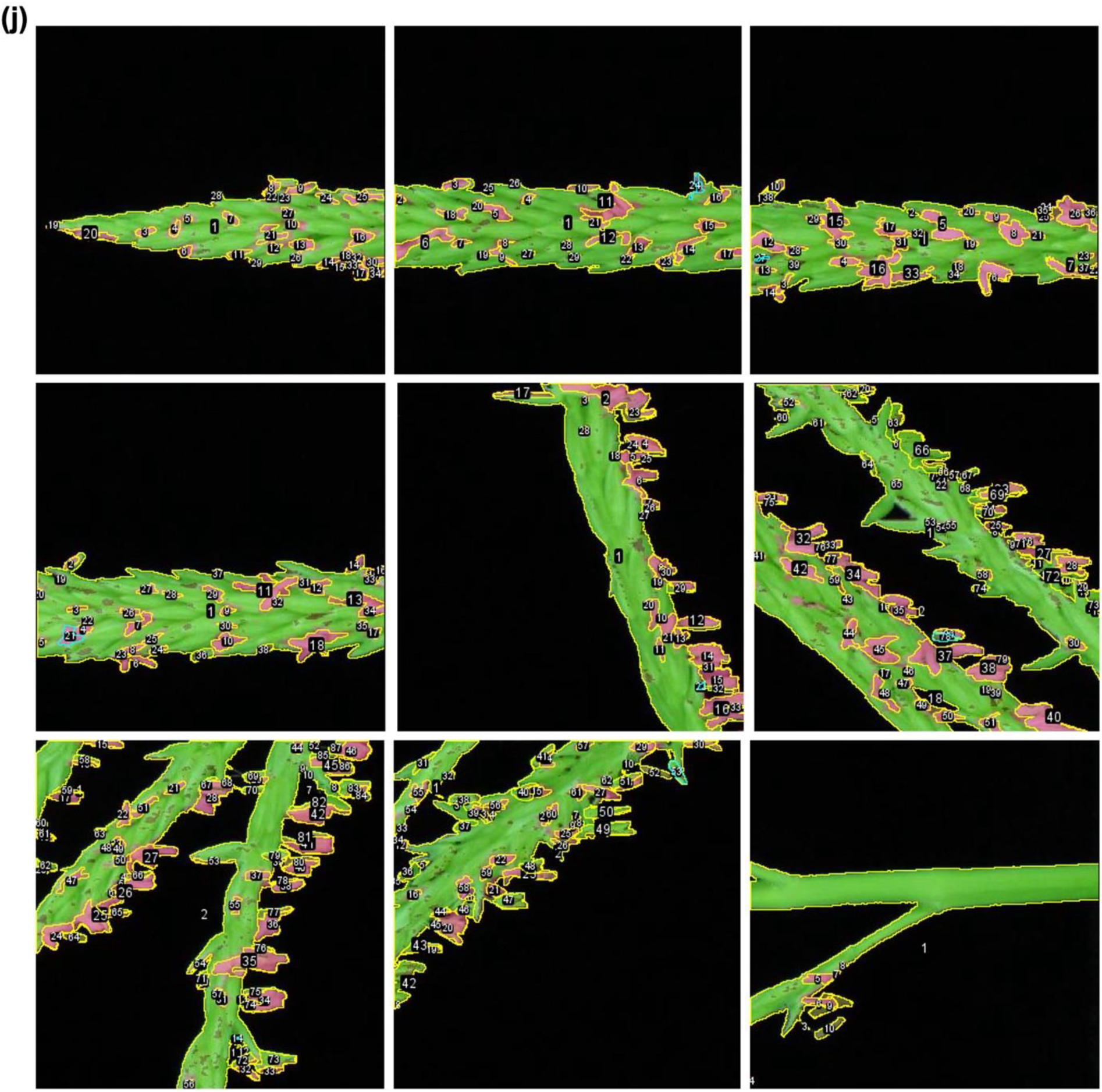
ROI and sums in processed tassel images for performance evaluation with PDF_1 for maize NAM founder line Oh43. TP of anther regions with fuchsia mask, FP of other tassel regions with fuchsia mask, FN of anther regions with green mask were manually labeled, annotated, and calculated with ImageJ. Sum of TP, FP, FN, and calculated recall, precision, and F_1_ scores are listed in Supporting Dataset.

**Supporting Figure 7k.**
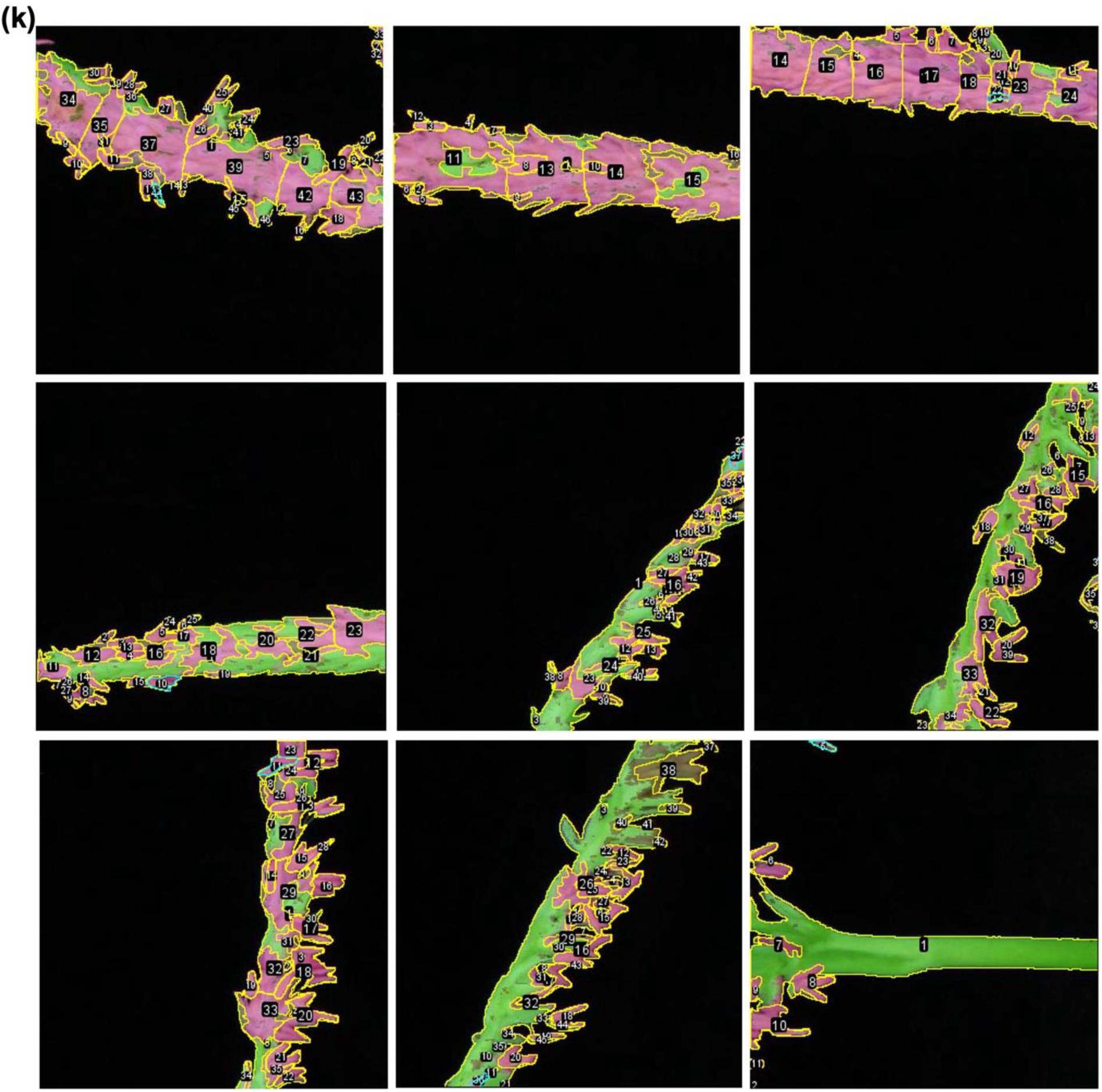
ROI and sums in processed tassel images for performance evaluation with PDF_1 for maize NAM founder line HP301. TP of anther regions with fuchsia mask, FP of other tassel regions with fuchsia mask, FN of anther regions with green mask were manually labeled, annotated, and calculated with ImageJ. Sum of TP, FP, FN, and calculated recall, precision, and F_1_ scores are listed in Supporting Dataset.

**Supporting Figure 7l.**
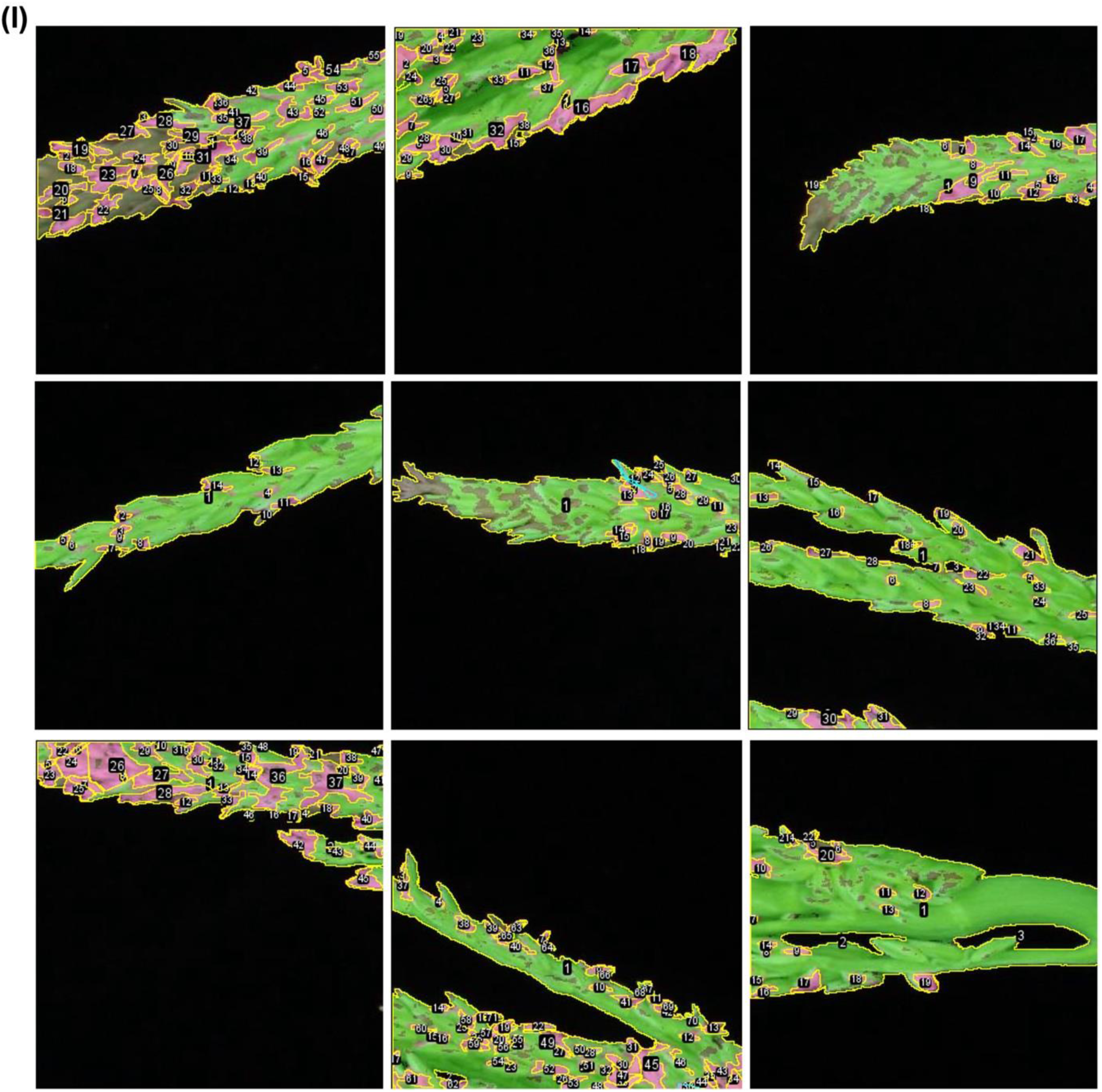
ROI and sums in processed tassel images for performance evaluation with PDF_1 for maize NAM founder line Tx303. TP of anther regions with fuchsia mask, FP of other tassel regions with fuchsia mask, FN of anther regions with green mask were manually labeled, annotated, and calculated with ImageJ. Sum of TP, FP, FN, and calculated recall, precision, and F_1_ scores are listed in Supporting Dataset.

**Supporting Figure 7m.**
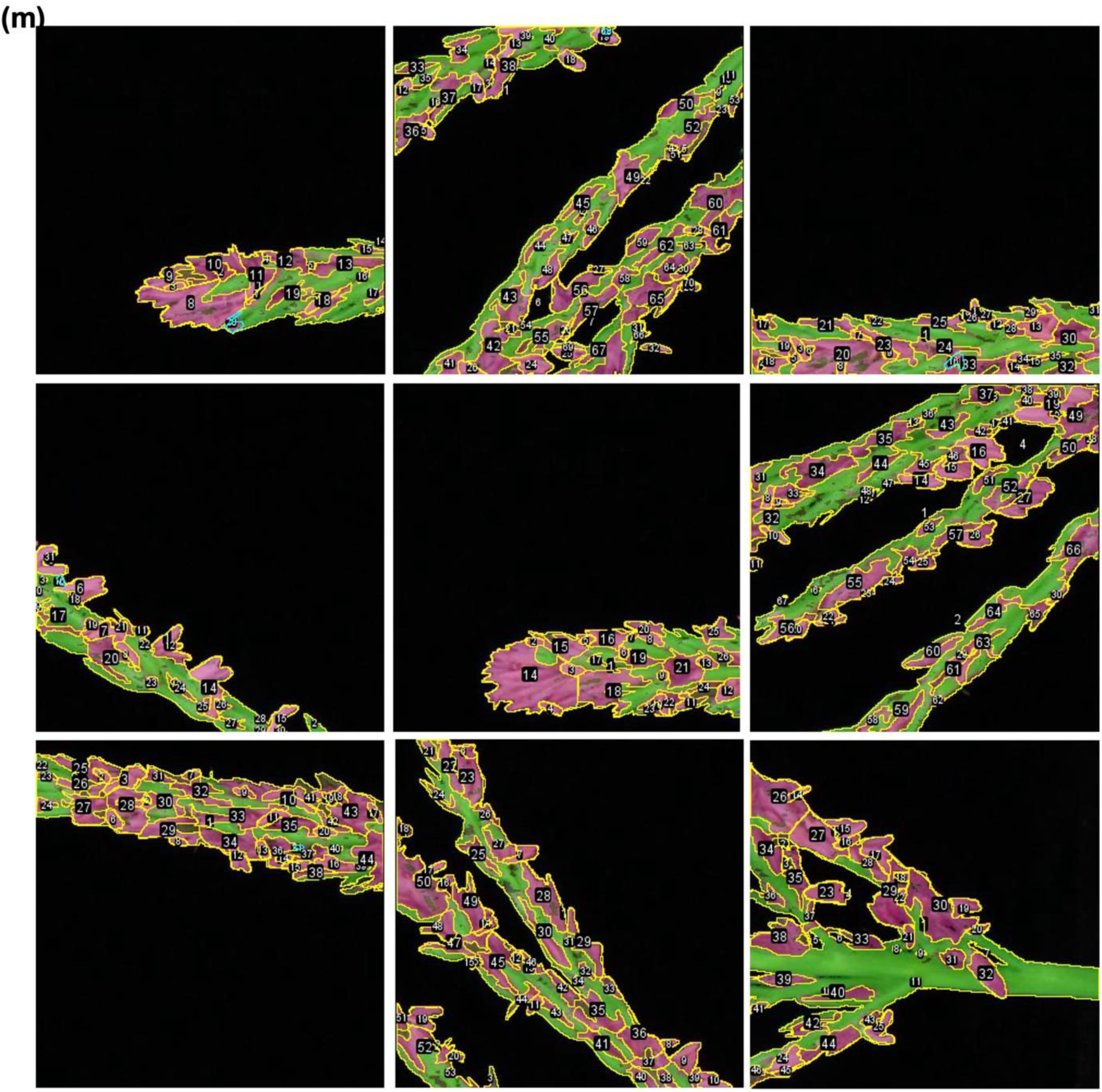
ROI and sums in processed tassel images for performance evaluation with PDF_1 for maize NAM founder line Ms71. TP of anther regions with fuchsia mask, FP of other tassel regions with fuchsia mask, FN of anther regions with green mask were manually labeled, annotated, and calculated with ImageJ. Sum of TP, FP, FN, and calculated recall, precision, and F_1_ scores are listed in Supporting Dataset.

**Supporting Figure 7n.**
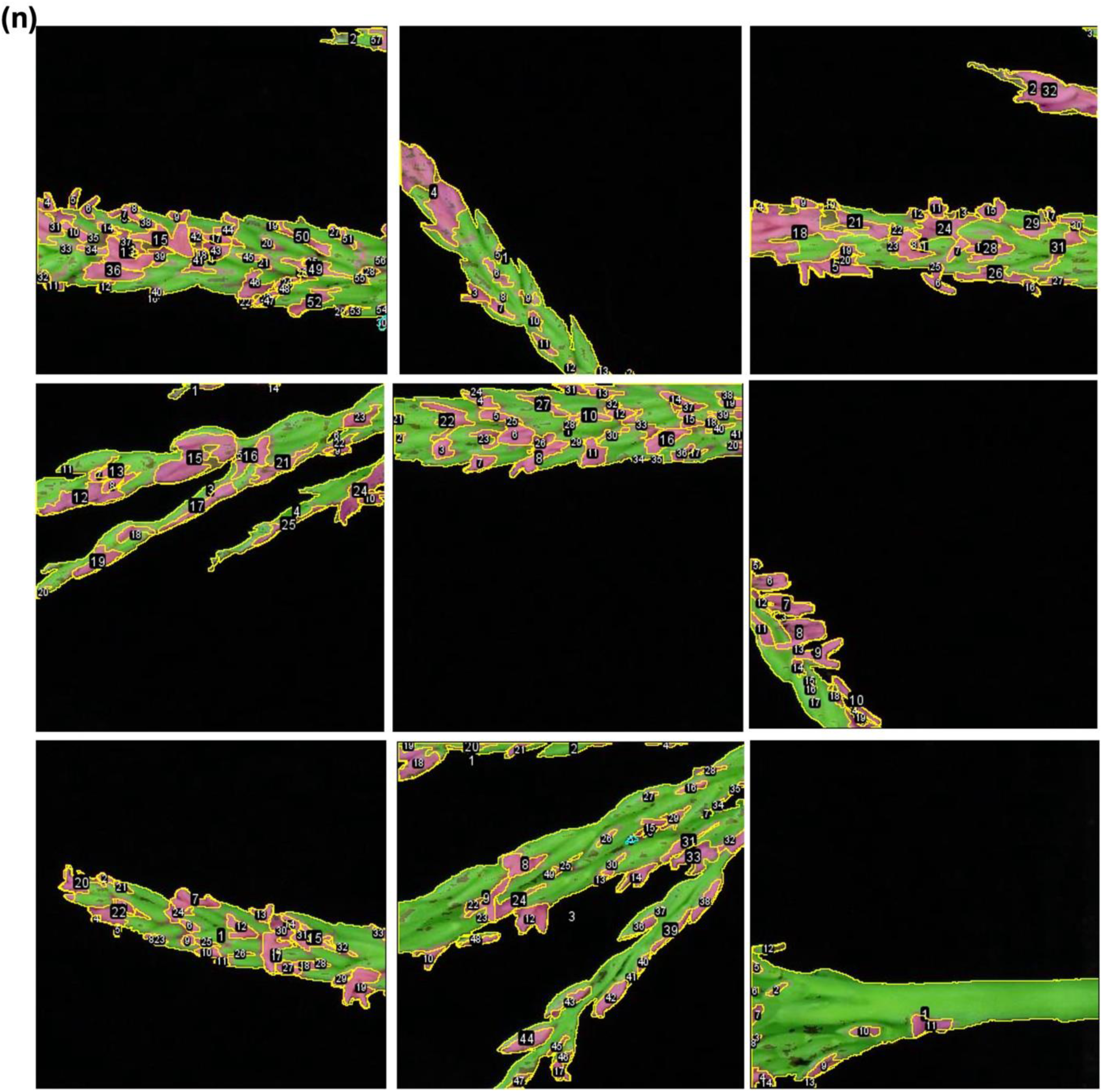
ROI and sums in processed tassel images for performance evaluation with PDF_1 for maize NAM founder line B97. TP of anther regions with fuchsia mask, FP of other tassel regions with fuchsia mask, FN of anther regions with green mask were manually labeled, annotated, and calculated with ImageJ. Sum of TP, FP, FN, and calculated recall, precision, and F_1_ scores are listed in Supporting Dataset.

**Supporting Figure 7o.**
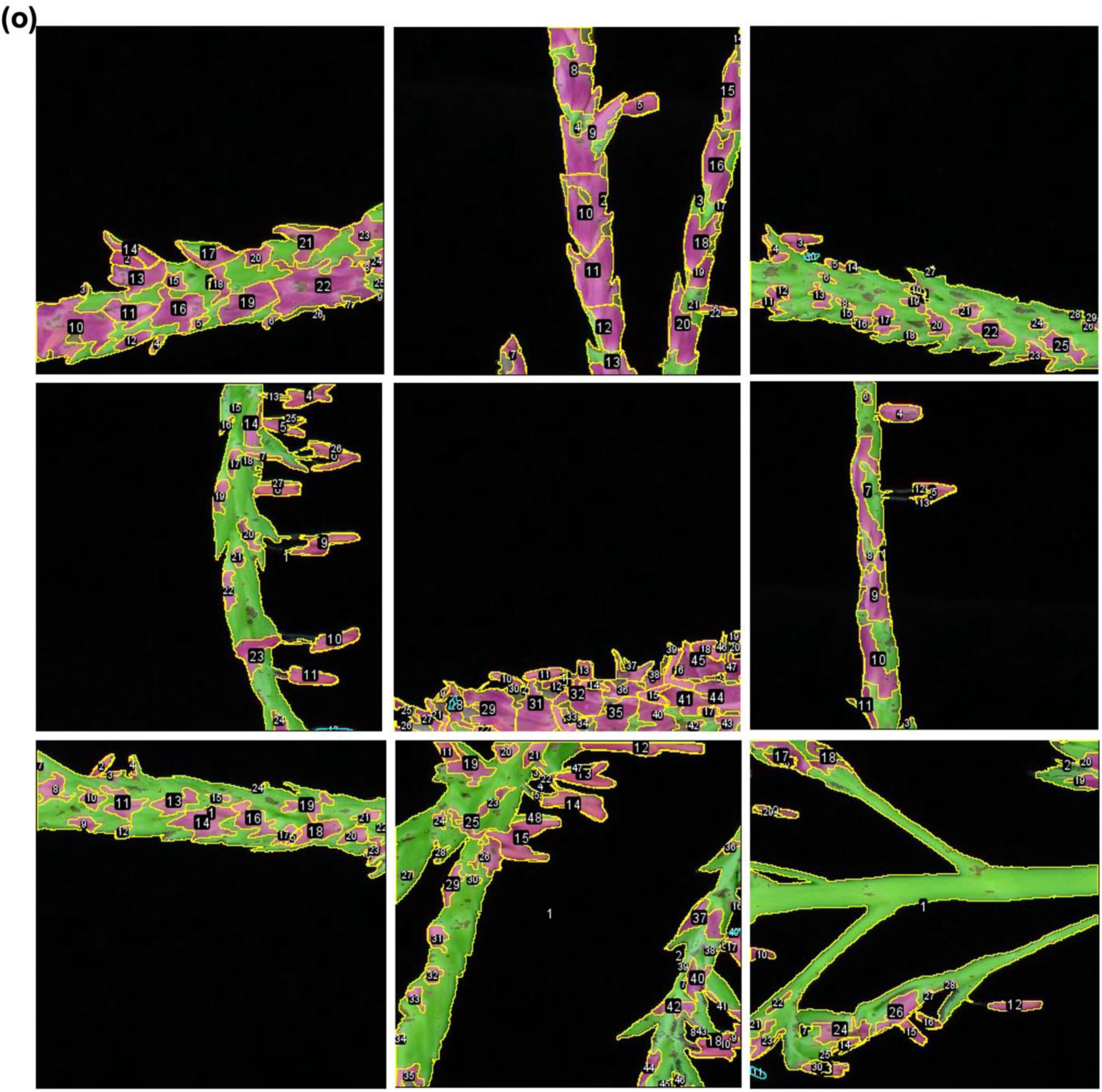
ROI and sums in processed tassel images for performance evaluation with PDF_1 for maize NAM founder line CML103. TP of anther regions with fuchsia mask, FP of other tassel regions with fuchsia mask, FN of anther regions with green mask were manually labeled, annotated, and calculated with ImageJ. Sum of TP, FP, FN, and calculated recall, precision, and F_1_ scores are listed in Supporting Dataset.

**Supporting Figure 7p.**
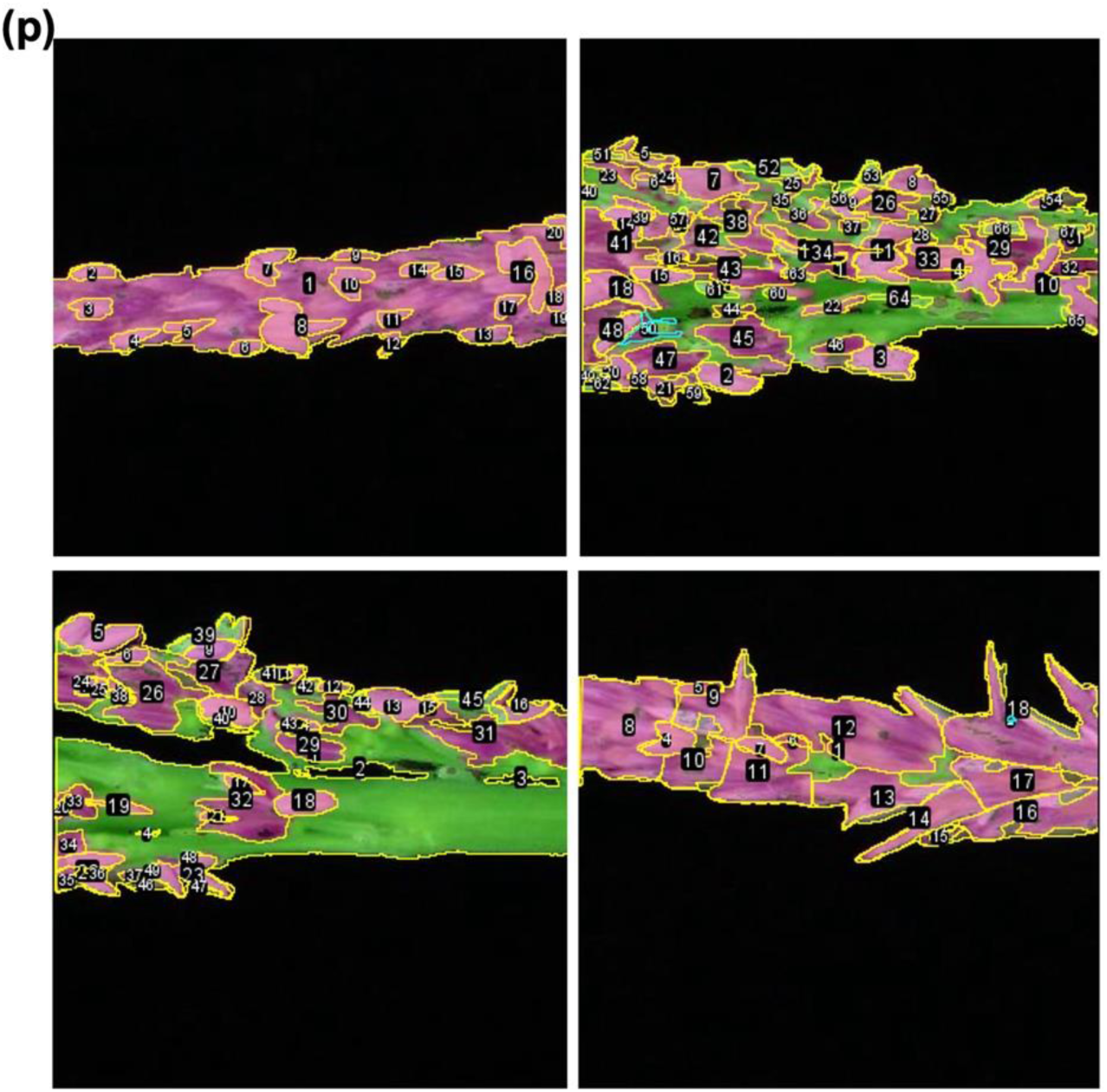
ROI and sums in processed tassel images for performance evaluation with PDF_1 for maize NAM founder line NC358. TP of anther regions with fuchsia mask, FP of other tassel regions with fuchsia mask, FN of anther regions with green mask were manually labeled, annotated, and calculated with ImageJ. Sum of TP, FP, FN, and calculated recall, precision, and F_1_ scores are listed in Supporting Dataset.

**Supporting Figure 7q.**
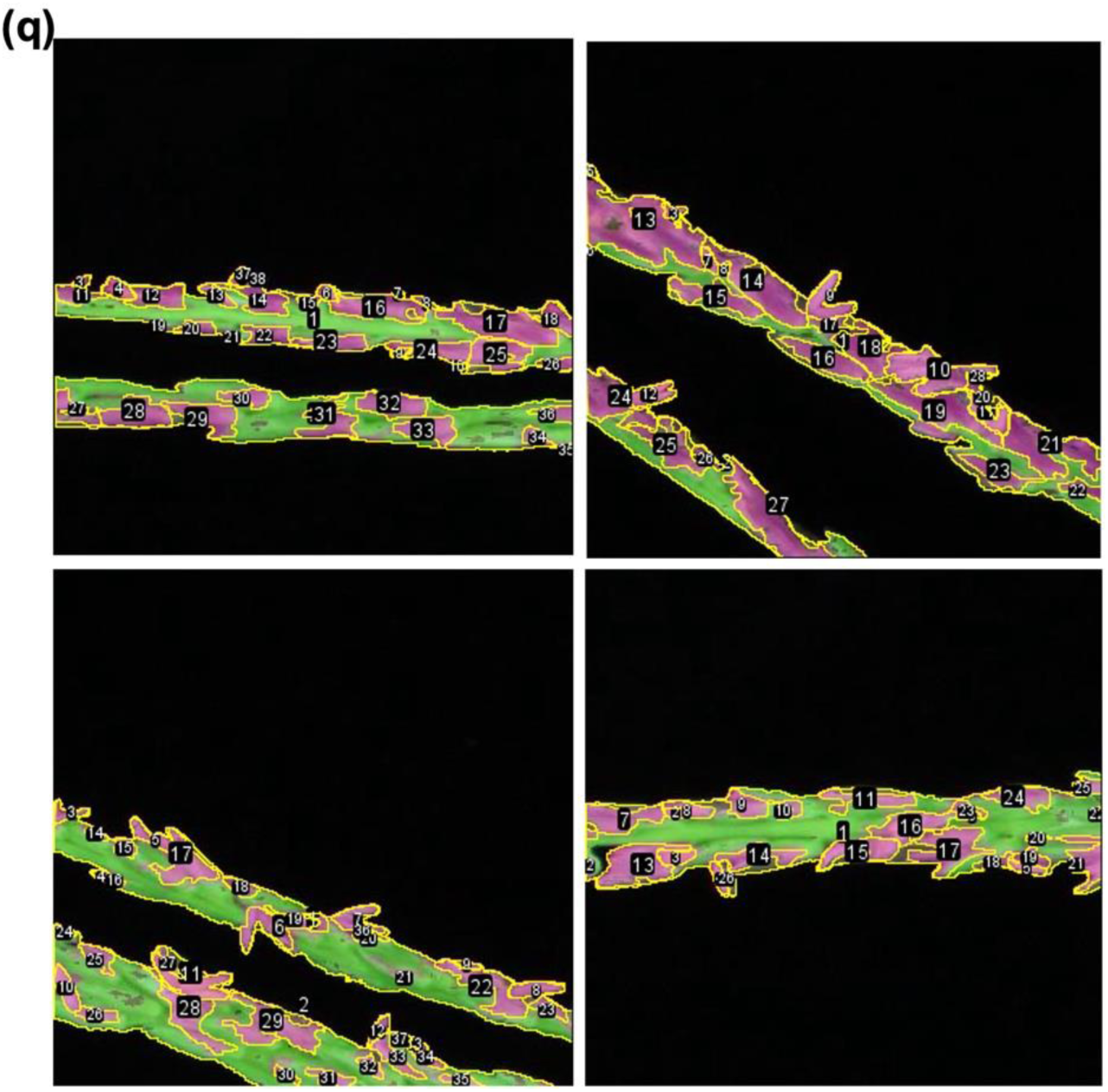
ROI and sums in processed tassel images for performance evaluation with PDF_1 for maize NAM founder line Ki11. TP of anther regions with fuchsia mask, FP of other tassel regions with fuchsia mask, FN of anther regions with green mask were manually labeled, annotated, and calculated with ImageJ. Sum of TP, FP, FN, and calculated recall, precision, and F_1_ scores are listed in Supporting Dataset.

**Supporting Figure 7r.**
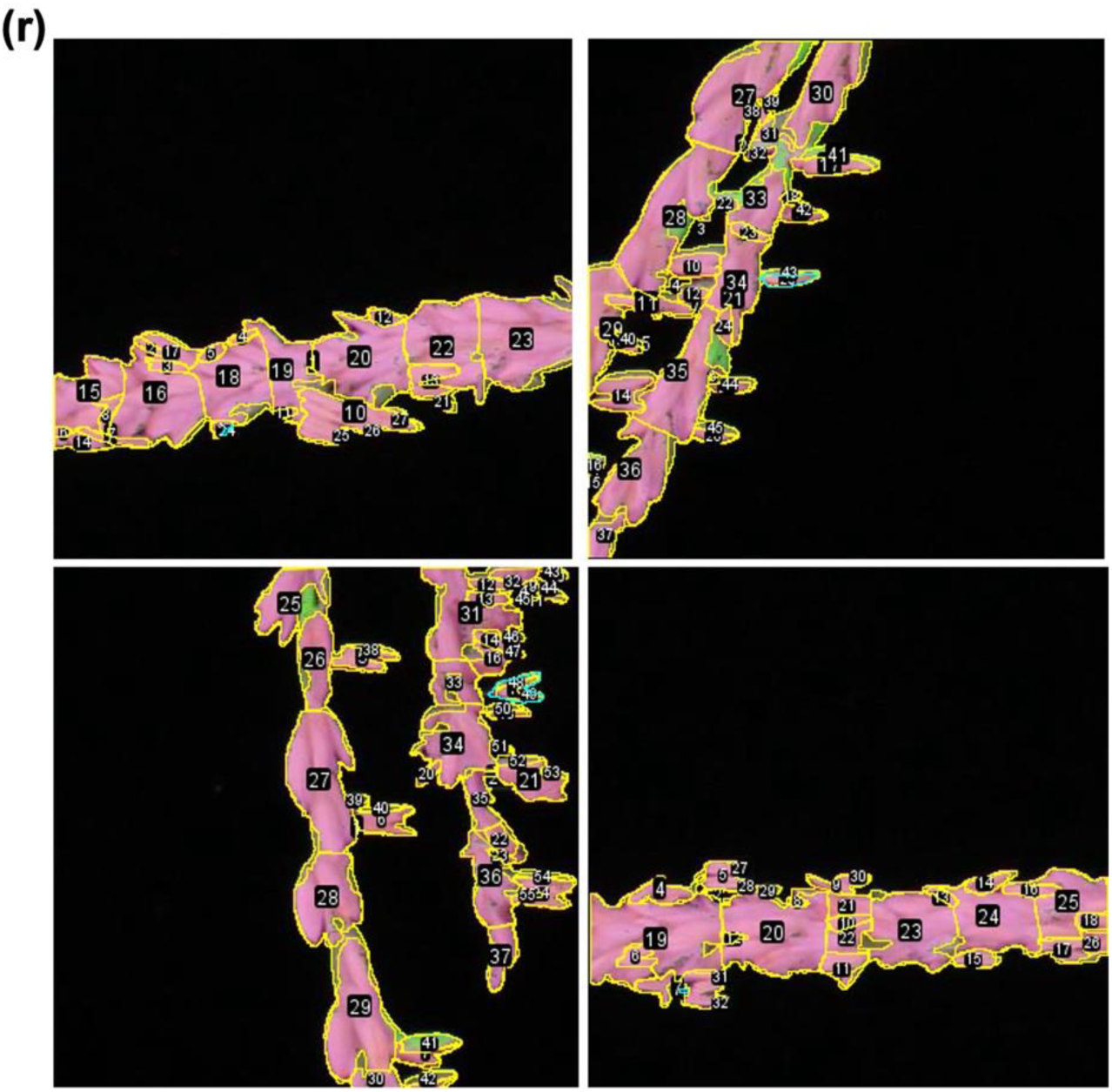
ROI and sums in processed tassel images for performance evaluation with PDF_1 for maize NAM founder line CML247. TP of anther regions with fuchsia mask, FP of other tassel regions with fuchsia mask, FN of anther regions with green mask were manually labeled, annotated, and calculated with ImageJ. Sum of TP, FP, FN, and calculated recall, precision, and F_1_ scores are listed in Supporting Dataset.

**Supporting Figure 7s.**
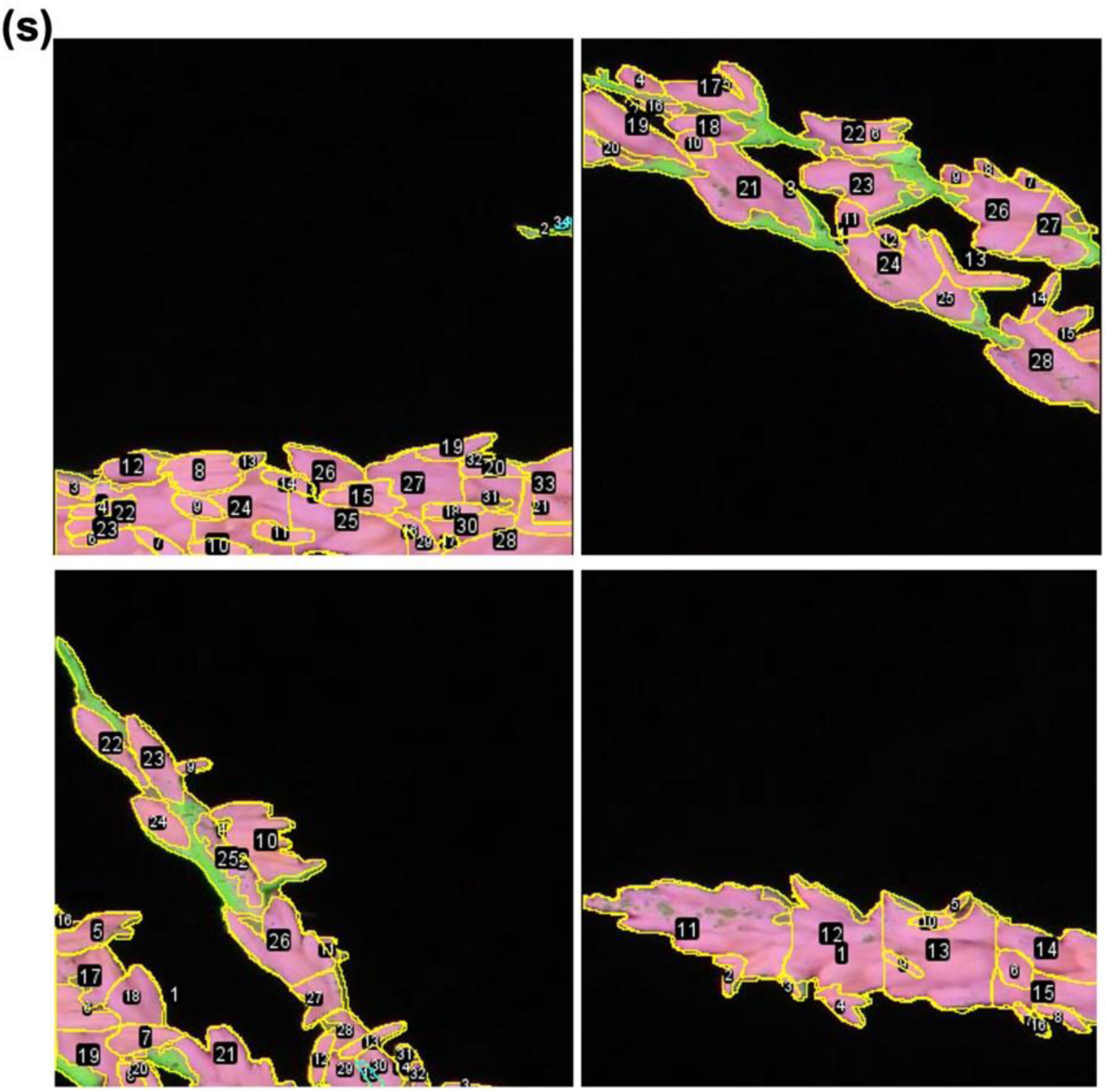
ROI and sums in processed tassel images for performance evaluation with PDF_1 for maize NAM founder line NC350. TP of anther regions with fuchsia mask, FP of other tassel regions with fuchsia mask, FN of anther regions with green mask were manually labeled, annotated, and calculated with ImageJ. Sum of TP, FP, FN, and calculated recall, precision, and F_1_ scores are listed in Supporting Dataset.

**Supporting Figure 7t.**
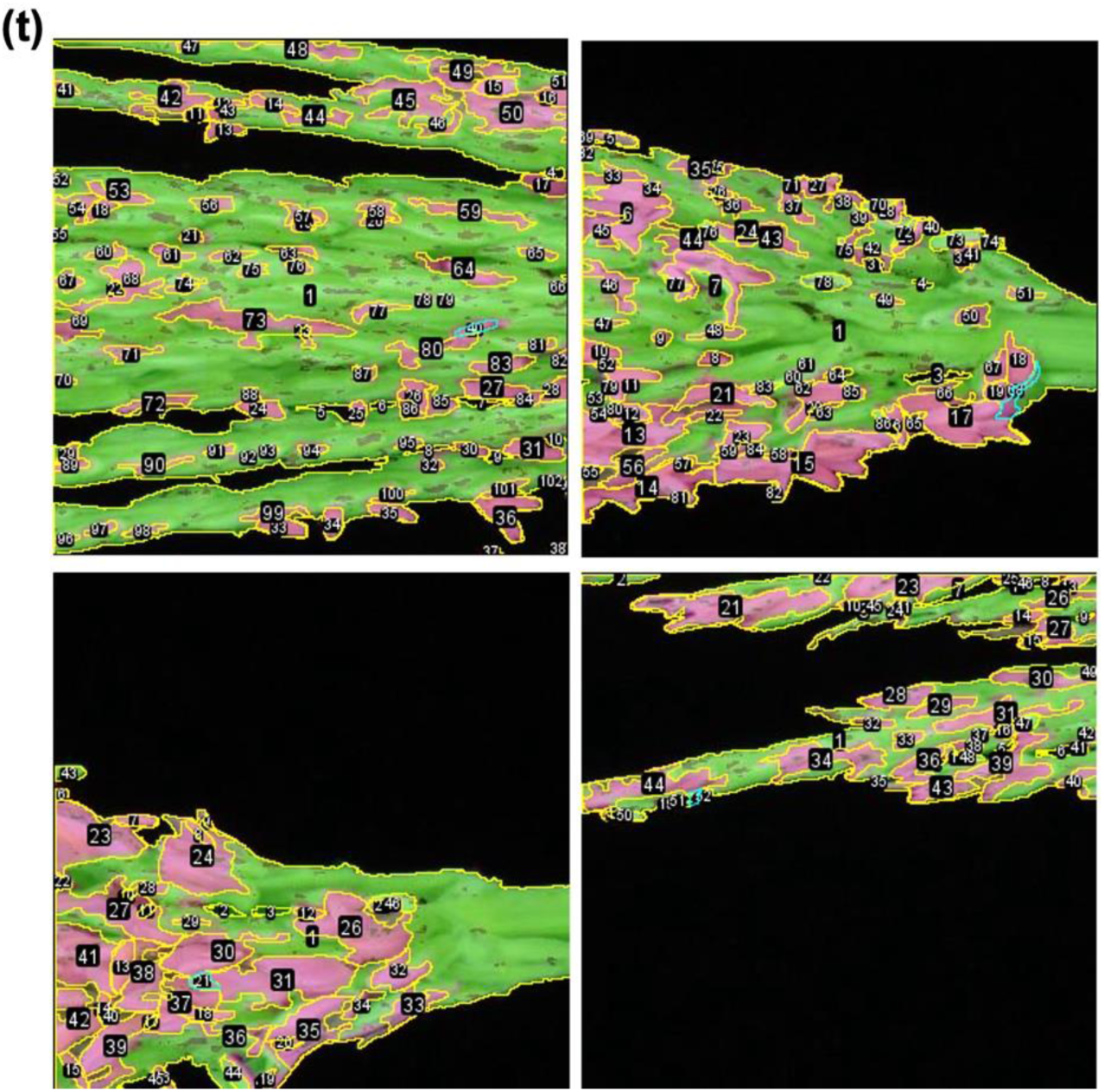
ROI and sums in processed tassel images for performance evaluation with PDF_1 for maize NAM founder line CML333. TP of anther regions with fuchsia mask, FP of other tassel regions with fuchsia mask, FN of anther regions with green mask were manually labeled, annotated, and calculated with ImageJ. Sum of TP, FP, FN, and calculated recall, precision, and F_1_ scores are listed in Supporting Dataset.

**Supporting Figure 7u.**
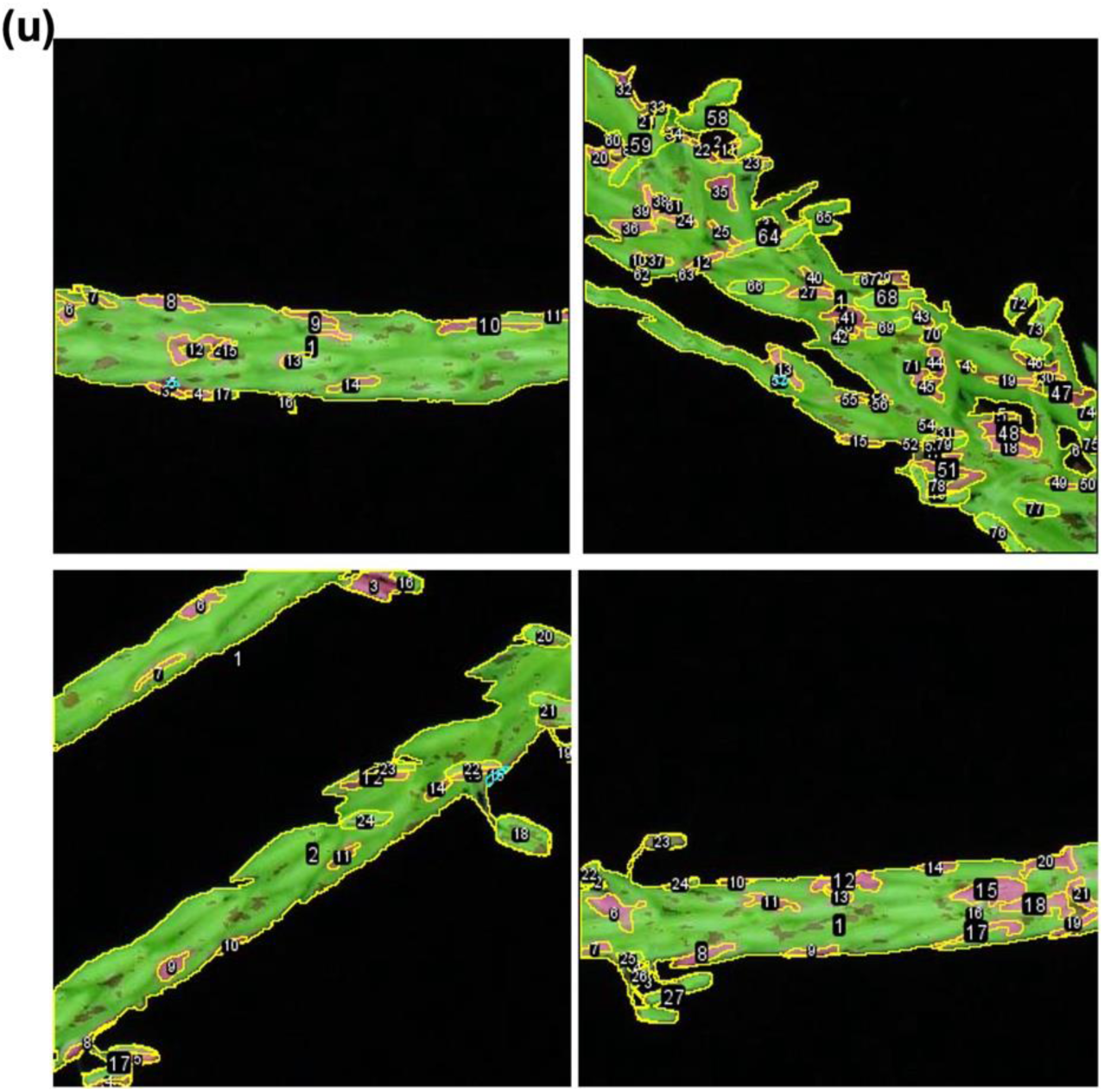
ROI and sums in processed tassel images for performance evaluation with PDF_1 for maize NAM founder line Mo17. TP of anther regions with fuchsia mask, FP of other tassel regions with fuchsia mask, FN of anther regions with green mask were manually labeled, annotated, and calculated with ImageJ. Sum of TP, FP, FN, and calculated recall, precision, and F_1_ scores are listed in Supporting Dataset.

**Supporting Figure 7v.**
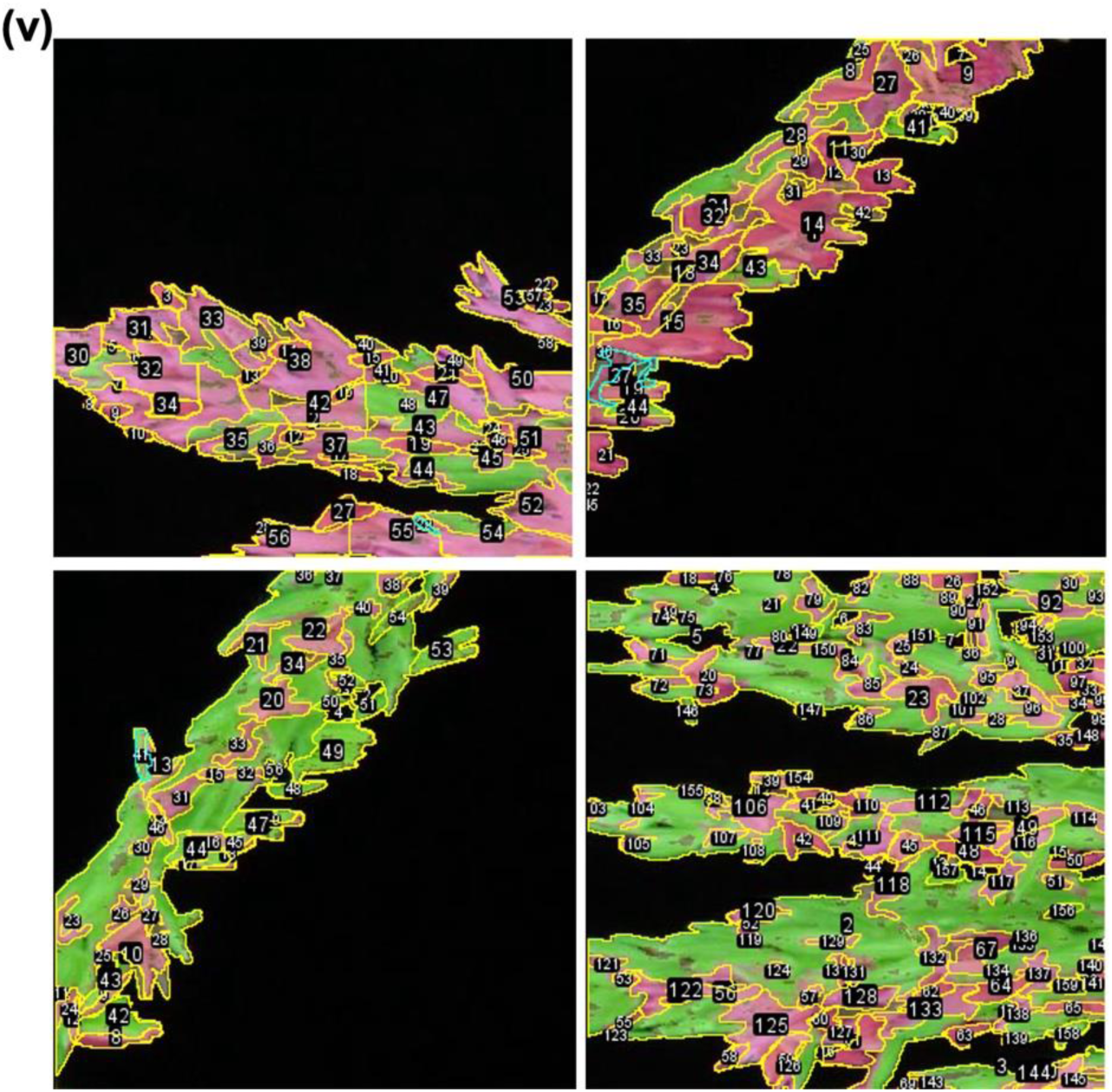
ROI and sums in processed tassel images for performance evaluation with PDF_1 for maize NAM founder line Ki3. TP of anther regions with fuchsia mask, FP of other tassel regions with fuchsia mask, FN of anther regions with green mask were manually labeled, annotated, and calculated with ImageJ. Sum of TP, FP, FN, and calculated recall, precision, and F_1_ scores are listed in Supporting Dataset.

**Supporting Figure 8.**
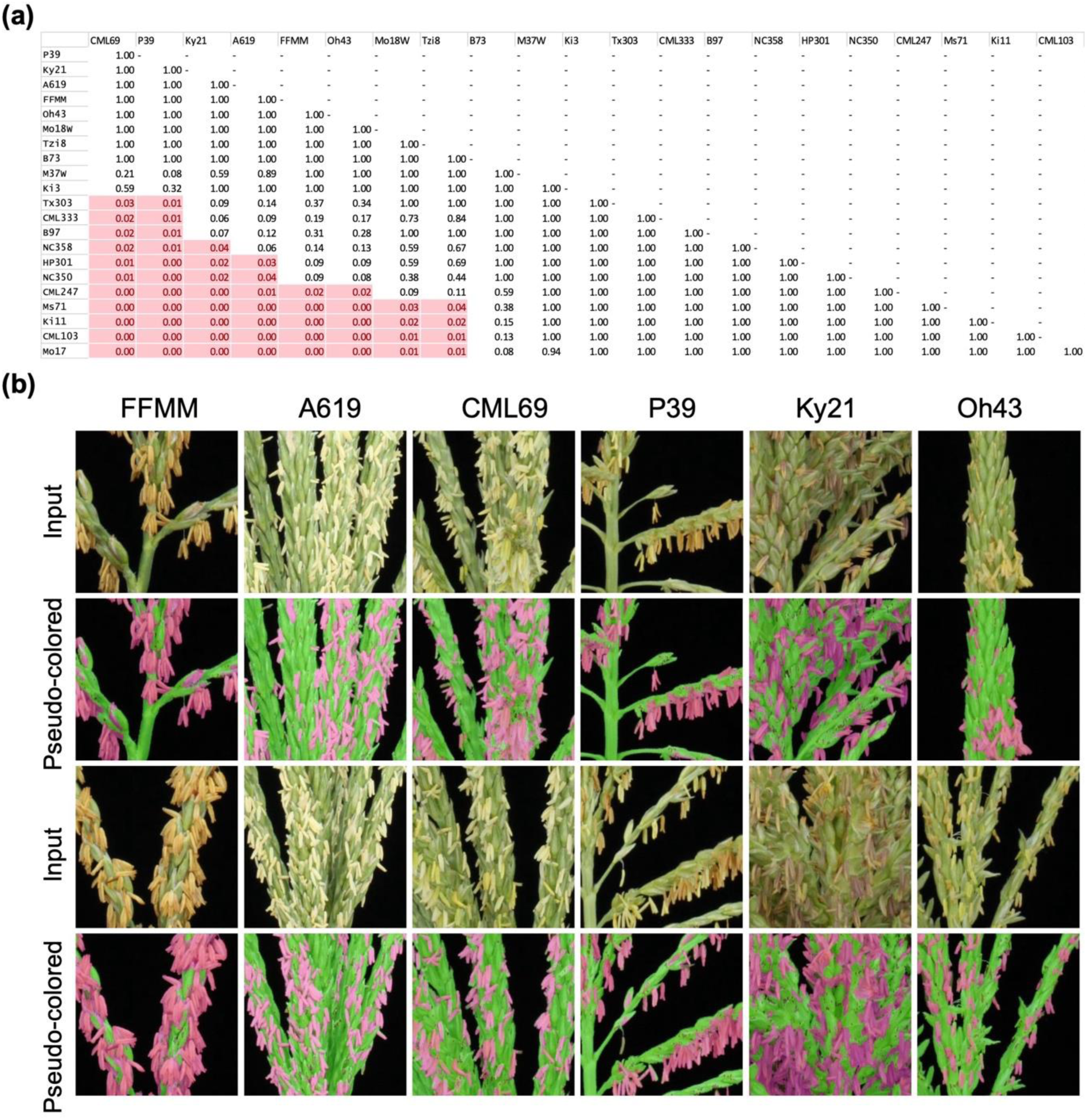
F_1_ score varies among maize lines. **(a)** Pairwise comparison of whole tassel anther ratios among different maize lines. The anther ratios of each group of plants from variable maize lines were tested for significance by one-way ANOVA and *post-hoc t*-test with Holm corrections. The pairs (horizontal - vertical) with significant differences (*p* < 0.05) highlighted in red. **(b)** Two more magnified example tassel input (upper) and pseudo-colored (lower) images of FFMM, A619, CML69, P39, Ky21, and Oh43.

**Supporting Figure 9.**
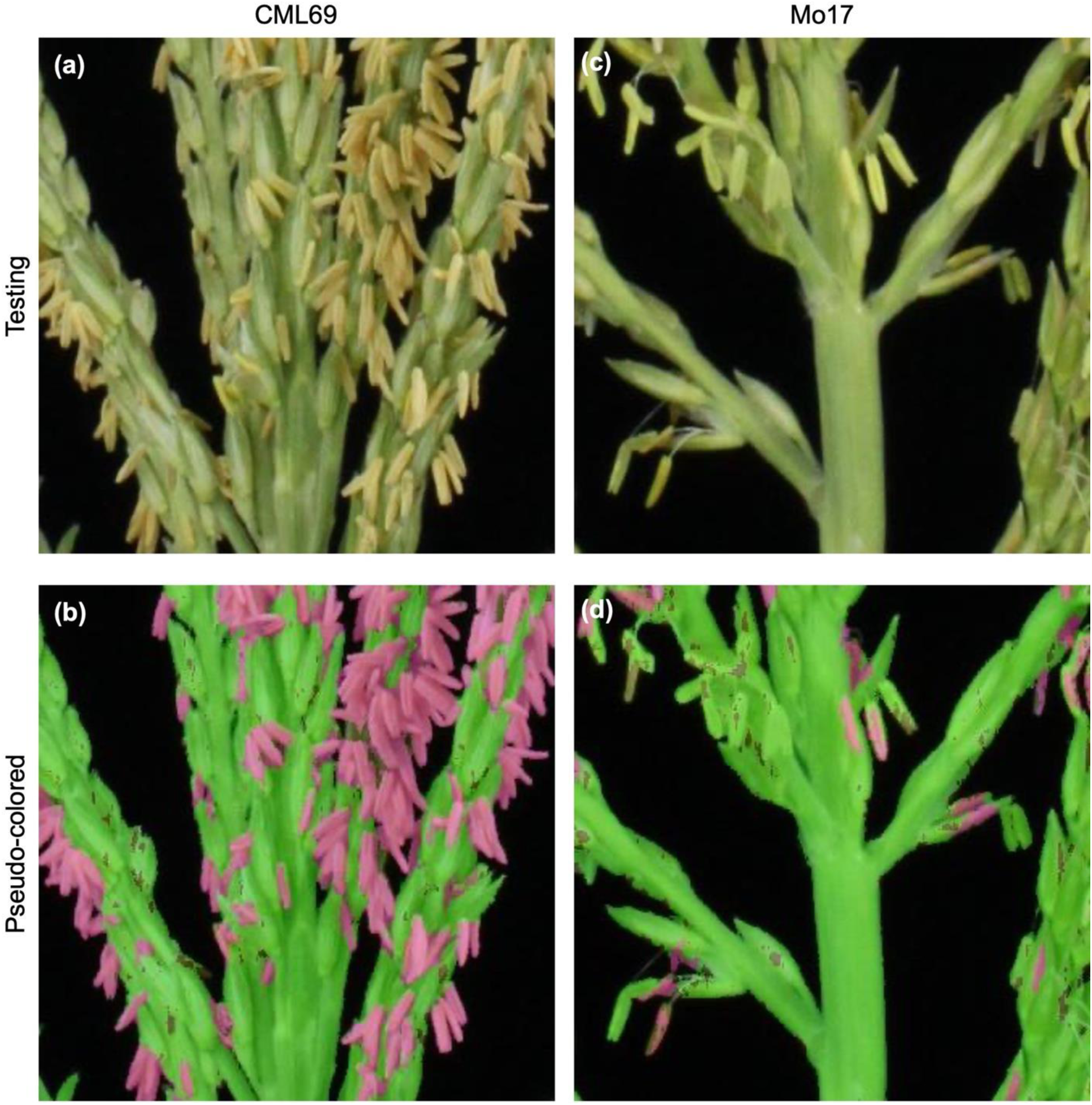
Comparison of CML69 with good segmentation and Mo17 with poor segmentation using PDF_1. Magnified original **(a)** and pseudo-colored **(b)** CML69 image. Anthers of CML69 in **(a)** are yellow and distinguishable from other tassel parts. This is not the case in magnified original image of Mo17 in **(c)**. Anthers of Mo17 are greenish, which resulted poor anther segmentation in **(d)**.

**Supporting Figure 10.**
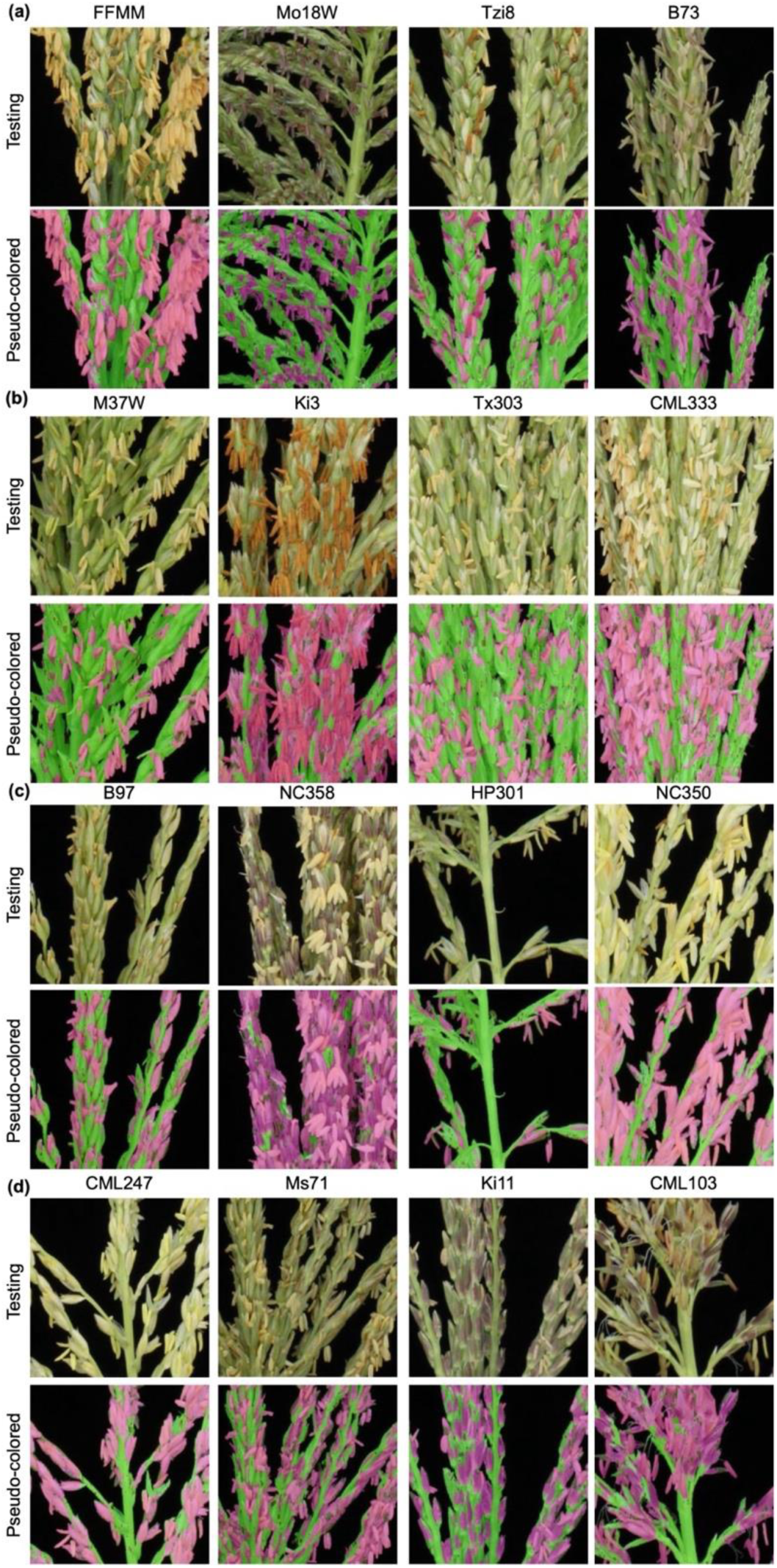
Performance of Tasselyzer varied due to different color patterns of maize lines. FFMM here is used as an example of good segmentation. Mo18W, Tzi8, B73, M37W, Ki3, Tx303 have some degrees of color patterns matching PDF_1, which allows Tasselyzer to perform average segmenting. CML333, B97, NC358, HP301, NC350, CML247, Ms71, Ki11, and CML103 have distinct color patterns which do not match PDF_1 to not allow Tasselyzer perform accurately.

**Supporting Figure 11.**
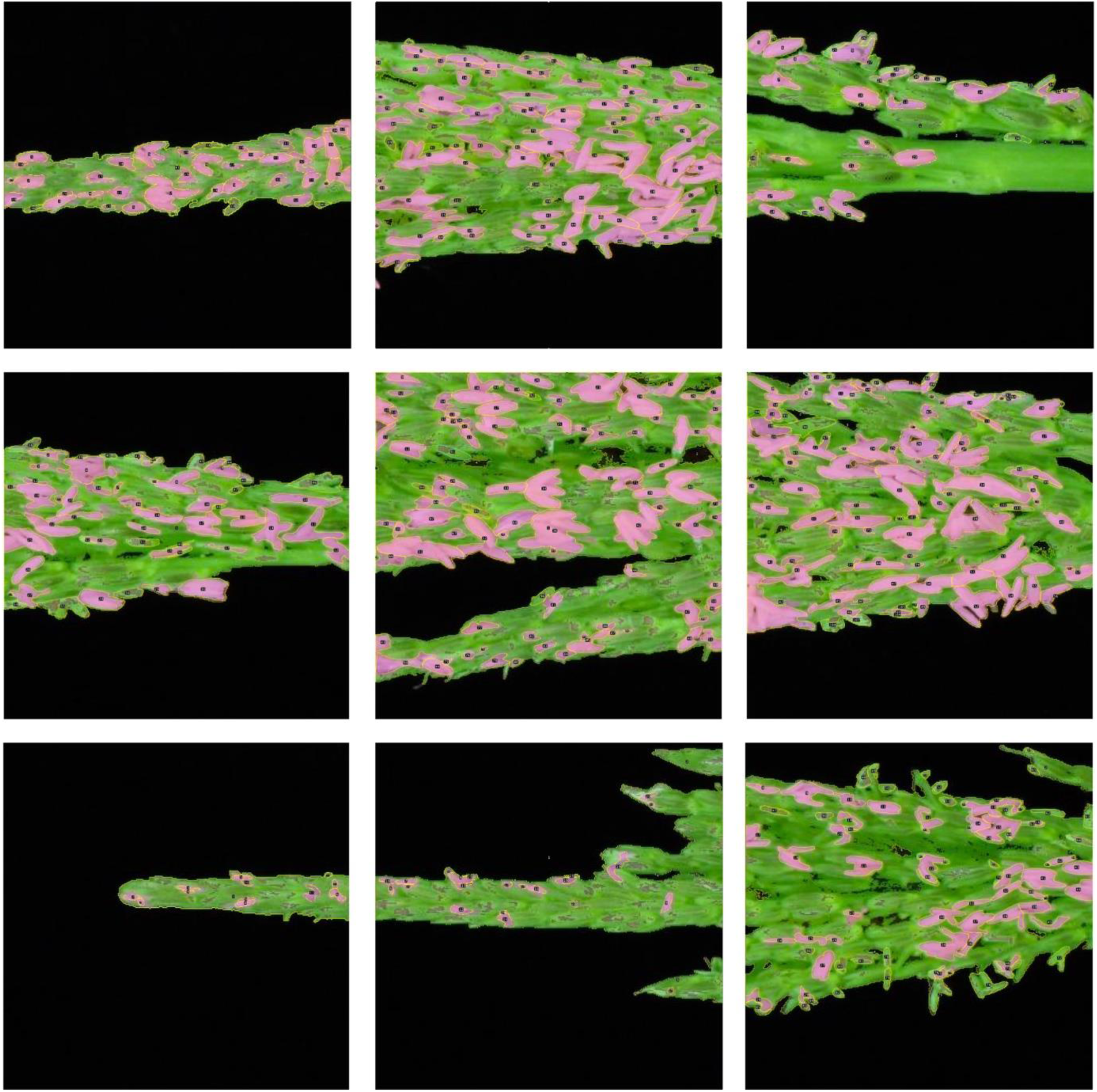
Regions of interest (ROI) and sums in processed tassel images for performance evaluation with PDF_2 for NC358. True positive (TP) of anther regions with fuchsia mask, false positive (FP) of other tassel regions with fuchsia mask, false negative (FN) of anther regions with green mask were manually labeled with “freehand selection” tool of ImageJ. Pseudo-colored images with yellow outlines of ROIs are demonstrated in the upper panels. Sum of TP, FP, FN, and calculated recall, precision, and F_1_ scores are listed in Supporting Dataset.

**Supporting Figure 12.**
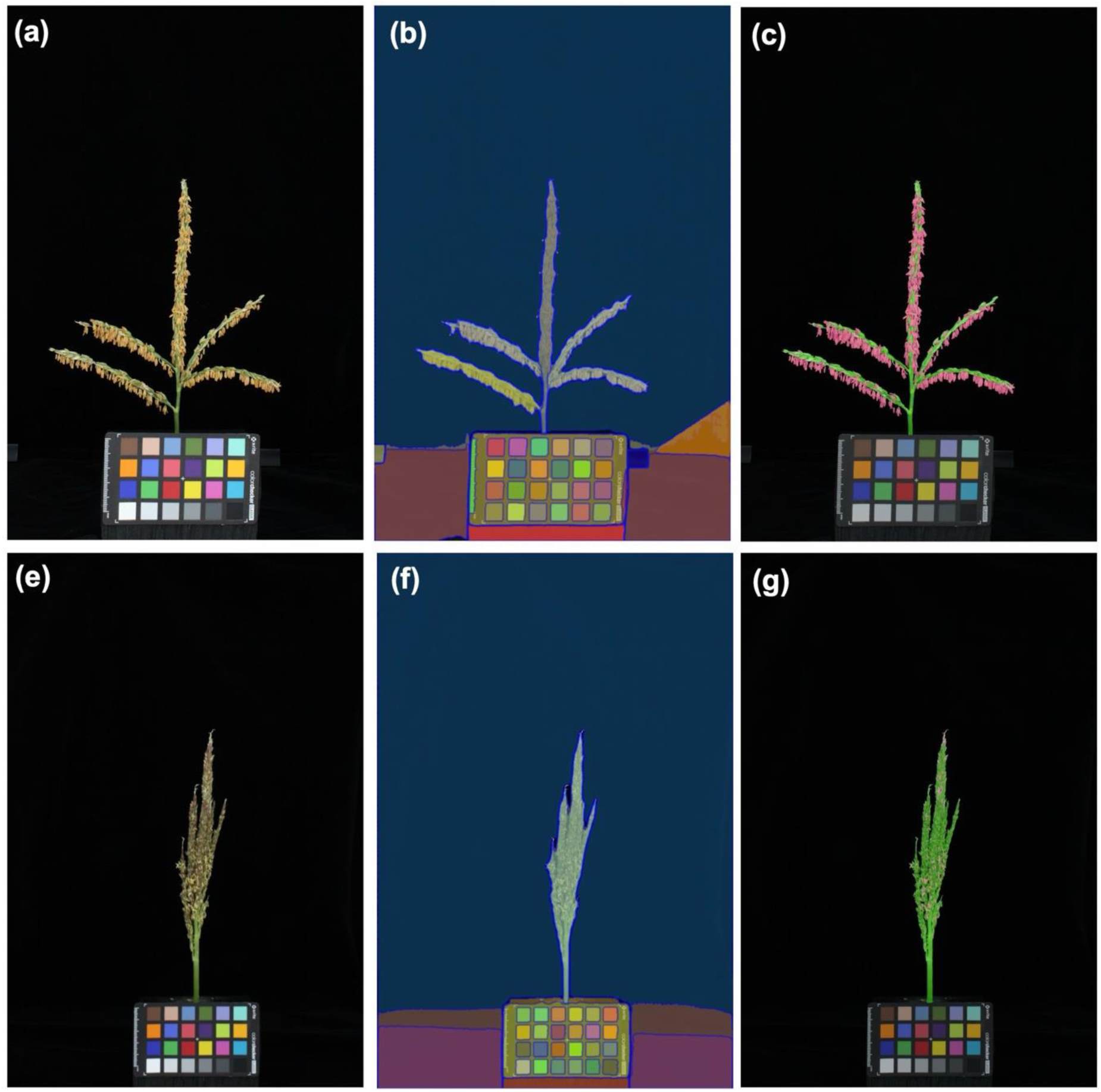
Performances of SAM on automated segmentation of tassels. An example of whole FFMM tassel image before **(a)** and after segmentation with the “everything” mode of SAM **(b)** and Tasselyzer **(c)**. An example of whole NC358 tassel image before **(e)** and after segmentation with everything mode of SAM **(f)** and Tasselyzer **(g)**. The everything mode of SAM allows automated segmentation unsupervised.

**Supporting Figure 13.**
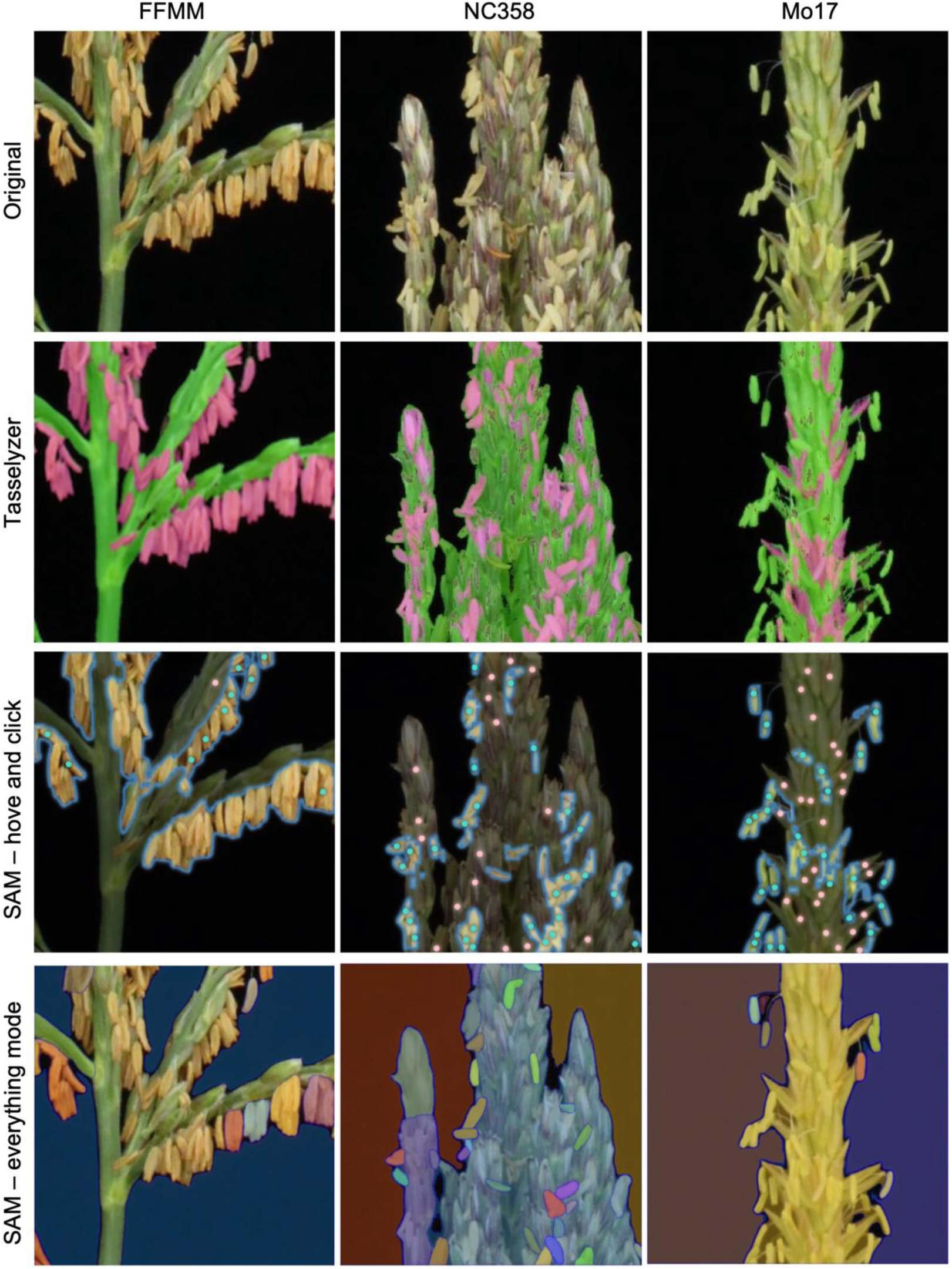
Performances of SAM on automated segmentation of partial tassel images of maize line FFMM, Mo17, and NC358. The manual “hover and click” and “everything” mode were employed, and the results are compared to those using Tasselyzer. The hover and click mode of SAM allows users to segment anthers by labeling anthers (blue dots) and exclude other tassel parts and background (pink dots) manually. The “everything mode” of SAM allows unsupervised automated segmentation.

**Supporting Table 1.** Summary of maize lines and images.

| Genotype | Type | Number of<br>Tassels | Number of<br>Images |
| --- | --- | --- | --- |
| B73<sup>a</sup> | NAM line, stiff corn | 5 | 20 |
| B97<sup>a</sup> | NAM line, non-stiff corn | 6 | 24 |
| Ky21<sup>a</sup> | NAM line, non-stiff corn | 7 | 16 |
| M37W<sup>a,b</sup> | NAM line, non-stiff corn | 6 | 25 |
| Mo17<sup>a</sup> | NAM line, non-stiff corn | 5 | 18 |
| Ms71<sup>a</sup> | NAM line, non-stiff corn | 7 | 23 |
| Oh43<sup>a</sup> | NAM line, non-stiff corn | 2 | 11 |
| CML103<sup>a</sup> | NAM line, tropical corn | 8 | 36 |
| CML247<sup>a</sup> | NAM line, tropical corn | 4 | 14 |
| CML69<sup>a,b</sup> | NAM line, tropical corn | 28 | 236 |
| CML333<sup>a</sup> | NAM line, tropical corn | 7 | 33 |
| Ki3<sup>a</sup> | NAM line, tropical corn | 6 | 44 |
| Ki11<sup>a</sup> | NAM line, tropical corn | 6 | 29 |
| NC350<sup>a</sup> | NAM line, tropical corn | 1 | 11 |
| Tx303<sup>a</sup> | NAM line, tropical corn | 4 | 21 |
| Tzi8<sup>a</sup> | NAM line, tropical corn | 9 | 51 |
| HP301<sup>a</sup> | NAM line, popcorn | 4 | 13 |
| P39<sup>a</sup> | NAM line, sweet corn | 2 | 14 |
| Mo18W<sup>a,b</sup> | Elite | 7 | 29 |
| NC358<sup>a</sup> | Elite | 5 | 31 |
| A619<sup>b,c</sup> | Inbred | 90 | 455 |
| FFMM<sup>b,c</sup> | FFMM-a, inbred | 75 | 288 |
| **Total** |  | 294 | 1,438 |

Notes:
<sup>a</sup> Research field grown in the summer of 2023, Missouri, USA.
<sup>b</sup> One image in this line was used for training for PDF\_1.
<sup>c</sup> Greenhouse grown in the summer of 2023 at Donald Danforth Plant Science Center, Missouri, USA.

**Supporting Table 2.** Summary of planting date and image numbers of A619 and FFMM.

| Group | Plant Date | Number of<br>A619<br>Tassels | Number of<br>A619<br>Images<sup>a</sup> | Number of<br>FFMM<br>Tassels | Number of<br>FFMM<br>Images<sup>b</sup> |
| --- | --- | --- | --- | --- | --- |
| Day 1 | 2023-7-2 | 7 | 35 | 6 | 27 |
| Day 2 | 2023-7-3 | 4 | 20 | 6 | 24 |
| Day 3 | 2023-7-4 | 4 | 25 | 7 | 26 |
| Day 4 | 2023-7-5 | 5 | 26 | 5 | 20 |
| Day 5 | 2023-7-6 | 8 | 40 | 6 | 25 |
| Day 6 | 2023-7-7 | 7 | 36 | 6 | 25 |
| Day 7 | 2023-7-8 | 8 | 41 | 6 | 30 |
| Day 8 | 2023-7-9 | 7 | 37 | 4 | 11 |
| Day 9 | 2023-7-10 | 6 | 30 | 5 | 22 |
| Day 10 | 2023-7-11 | 6 | 30 | 6 | 18 |
| Day 11 | 2023-7-12 | 7 | 35 | 5 | 19 |
| Day 12 | 2023-7-13 | 7 | 35 | 5 | 20 |
| Day 13 | 2023-7-14 | 5 | 26 | 4 | 11 |
| Day 14 | 2023-7-15 | 8 | 39 | 4 | 10 |
| **Total** |  | 89 | 455 | 75 | 288 |

Notes:
<sup>a</sup> All tassels of A619 were detached and imaged freshly on 2023-8-29.<sup>b</sup> All tassels of FFMM were detached and imaged freshly on 2023-8-12.

